# Systematic Evaluation of Normalization Approaches in Tandem Mass Tag and Label-Free Protein Quantification Data Using PRONE

**DOI:** 10.1101/2025.01.27.634993

**Authors:** Lis Arend, Klaudia Adamowicz, Johannes R. Schmidt, Yuliya Burankova, Olga Zolotareva, Olga Tsoy, Josch K. Pauling, Stefan Kalkhof, Jan Baumbach, Markus List, Tanja Laske

## Abstract

Despite the significant progress in accuracy and reliability in mass spectrometry technology, as well as the development of strategies based on isotopic labeling or internal standards in recent decades, systematic biases originating from non-biological factors remain a significant challenge in data analysis. In addition, the wide range of available normalization methods renders the choice of a suitable normalization method challenging. We systematically evaluated 17 normalization and two batch effect correction methods, originally developed for pre-processing DNA microarray data but widely applied in proteomics, on six publicly available spike-in and three label-free and tandem mass tag datasets. Opposed to state-of-the-art normalization practice, we found that a reduction in intragroup variation is not directly related to the effectiveness of the normalization methods. Furthermore, our results demonstrated that the methods RobNorm and Normics, specifically developed for proteomics data, in line with LoessF performed consistently well across the spike-in datasets, while EigenMS exhibited a high false positive rate. Finally, based on experimental data, we show that normalization substantially impacts downstream analyses, and the impact is highly dataset-specific, emphasizing the importance of use-case-specific evaluations for novel proteomics datasets. For this, we developed the PROteomics Normalization Evaluator (PRONE), a unifying R package enabling comparative evaluation of normalization methods, including their impact on downstream analyses, while offering considerable flexibility, acknowledging the lack of universally accepted standards. PRONE is available on Bioconductor with a web application accessible at https://exbio.wzw.tum.de/prone/.

## Introduction

High-throughput omics technologies nowadays produce massive amounts of data and steadily progress in detection accuracy and data generation speed (1). However, non-biological factors arising from variations in biological experiments, sample preparation, instrumental analyses and raw data processing, frequently introduce systematic biases during an experiment (**Figure 1A**) (2,3). Failure to address these biases can lead to erroneous results and misleading conclusions in downstream analyses, such as differential expression (DE) and functional enrichment analysis (2,4,5). Given that achieving perfect experimental precision in the lab is nearly impossible, the critical step in compensating for this experimental variability is data normalization (1,3). Data normalization aims to remove or minimize these systematic biases while preserving the biological signal of interest. However, the nature and magnitude of the bias in the data is typically unknown beforehand since it is not easy to measure nor to quantify, thus the choice of a suitable approach among the wide range of available normalization methods poses a notable challenge (2,3,6).

**Figure 1:**
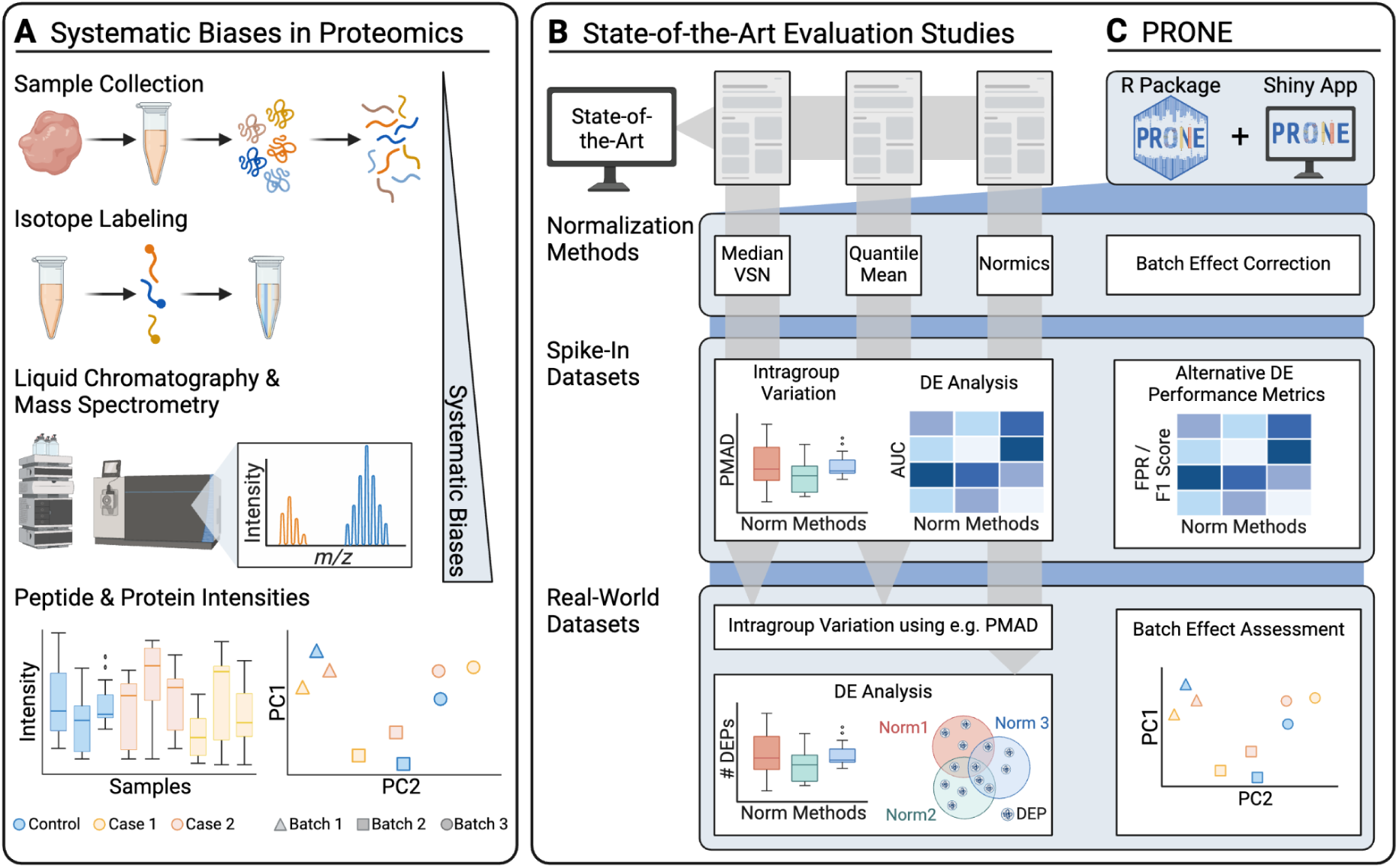
O**v**erview **of the evaluation study.** (A) MS-based proteomics datasets are affected by systematic biases, attributed to variations in experiment conditions during MS analysis, ranging from sample handling to differences introduced by the instrumentation. These biases distort sample distributions and introduce potential batch effects, particularly in TMT datasets. Existing evaluation studies on proteomics normalization focus on distinct sets of normalization methods, assess intragroup variation on spike-in datasets, and predominantly use the area under the receiver operating characteristic curve (AUC) to analyze the differential expression (DE) results for each normalization method. Most studies conclude their evaluation on biological datasets after calculating intragroup variation metrics due to the absence of a known ground truth. Only very few studies proceed with examining DE results on biological datasets. To perform our comprehensive benchmarking, we developed PRONE, which integrates a broad array of normalization methods and provides batch effect correction, PRONE also expands on previous studies by calculating alternative performance metrics, including false positive rate and F1 score, rather than relying solely on AUC values. Additionally, batch effects can be evaluated using multiple techniques within the PRONE framework.

The majority of existing evaluation studies (2,4–6) and graphical interfaces, such as Normalyzer (1), NormalyzerDE (7), and proteiNorm (8), have been developed to systematically evaluate normalization methods, such as median centering, quantile normalization, linear and local regression normalization, and variance stabilization normalization (VSN) for label-free quantification (LFQ) (**Figure 1B**). These studies often evaluate a diverse set of normalization methods on protein-level proteomics data, with many normalization methods originating from the DNA microarray analysis techniques. Newer approaches such as RobNorm (9) and Normics (10), specifically designed for proteomics applications, have yet to be subjected to independent validation. Importantly, most evaluations are limited in assessing intragroup variation on experimental biological datasets without known ground truth and do not extend to evaluating the impact of normalization on downstream analyses, such as DE analysis, due to the lack of a known ground truth (1,2,4,7,8). Nevertheless, the impact of normalization on the identification and quantity of differentially abundant proteins between sample groups can be analyzed and was observed in a limited number of studies (6,9,10), yet this capability has not been incorporated as a feature in any tool. Furthermore, most studies restrict the evaluation on LFQ datasets. However, tandem mass tag (TMT), a chemical labeling method that enables the simultaneous MS-analysis of up to 18 samples pooled together, has become state-of-the-art in large-scale proteomic studies (11,12). The integration of multiple TMT batches within a single analysis introduces batch effects and thus reduces data quality. While optimized experimental design set-ups are considered to minimize batch effects (13,14), further computational methods for batch effect correction, such as internal reference scaling (IRS) (15), are required.

While the existing studies and graphical interfaces show that a comprehensive comparative evaluation is indispensable in identifying an appropriate normalization strategy for a specific dataset, their limitations, such as the exclusion of downstream analysis evaluation and the lack of focus on TMT datasets, remain apparent. Building upon these limitations, our study aims to address the gaps by providing a systematic evaluation of a comprehensive literature-based collection of 17 normalization strategies and two batch effect correction methods commonly employed in proteomics (**Figure 1C**, see **Methods**). We utilized six biological proteomics datasets with known ground truth, referred to as spike-in datasets, allowing for ranking the methods in their ability to detect expected differences between sample groups. Additionally, we applied the normalization techniques to one LFQ and two TMT biological proteomics datasets, which are absent of spike-in proteins and are referred to as biological datasets, to compare their performance in a typical real research scenario without known ground truth.

To perform this comprehensive benchmarking and streamline the selection process of optimal normalization techniques for other proteomics data, we developed the Proteomics Normalization Evaluator (PRONE), an R package available on Bioconductor with an accompanying graphical interface accessible at https://exbio.wzw.tum.de/prone/ (**Figure 1C**). PRONE incorporates 17 normalization techniques and two batch effect correction methods in line with multiple quantitative and qualitative functions to evaluate the performance of the methods. In addition, PRONE offers an integrated solution for the complete analysis of an LFQ or TMT proteomics dataset, including missing value handling, outlier detection, and DE analysis. Unlike other interfaces that allow DE analysis based on a single selected normalization technique, PRONE enables the comparative evaluation of DE results across multiple normalization methods, highlighting the impact of normalization on downstream analyses. It thus provides a robust and reliable framework for proteomics data analysis without the need to install or integrate multiple separate packages.

## Materials and methods

### Data description

#### Spike-in datasets

The experimental design of spike-in datasets, which involves adding specific proteins at known concentrations to a constant background proteome, such as yeast lysates or human cell extracts, allows for evaluating the normalization methods on their ability to detect DE and calculating performance metrics. We obtained six publicly available spike-in datasets (**Table 1** and **Supplementary Methods**): two protein standard (UPS1) spike-in (dB1-dB2, 16,17), three *Escherichia coli* spike-in (dB3-dB5, 18–20), and one yeast spike-in dataset (dB6, 21).

**Table 1:** Spike-in and biological datasets. Spike-in datasets are denoted with the prefix “dS,” while biological datasets without a known ground truth are consistently denoted with “dB.” In the column for quantification type, brackets indicate the number of batches used for the specific TMT dataset. The identifiers from ProteomeXchange.org are added for the raw data column, except for dB2, where the PDC study identifier is utilized. Due to the inconsistency in the availability of raw data from the original studies, quantification data was sourced from various other studies that have released the protein intensity matrix. Further descriptions, including the utilization of instrumentation, are provided in the Supplementary Methods.

| Dataset | Type | Quantification Type | Raw Data (ID) | Quantification Data |
| --- | --- | --- | --- | --- |
| dS1 | UPS1 spike-in<br>(4 levels) | LFQ | Tabb *et al.* (16)<br>(CPTAC Study 6) | Välikangas *et al.* (2) |
| dS2 | UPS1 spike-in<br>(6 levels) | LFQ | Ramus *et al.* (17)<br>(PXD001819) | Graw *et al.* (8) |
| dS3 | *E.coli* spike-in<br>(5 levels) | LFQ | Shen *et al.* (18)<br>(PXD003881) | Sticker *et al.* (25) |
| dS4 | *E.coli* spike-in<br>(2 levels) | LFQ | Cox *et al.* (19)<br>(PXD00279) | Cox *et al.* (19)<br>(PXD00279) |
| dS5 | *E.coli* spike-in | TMT 10-plex (1) | Zhu *et al.* (20) | Phil Wilmarth (26) |
|  | (3 levels) |  | (PXD013277) |  |
| dS6 | yeast spike-in<br>(3 levels) | TMT 11-plex (1) | O'Connell *et al.* (21)<br>(PXD007683) | Ammar *et al.* (27) |
| dB1 | Osteogenic differentiation of<br>hPDLSCs (4 time points) | TMT 6-plex (3) | Li *et al.* (22)<br>(PXD020908) | MaxQuant executed<br>in-house |
| dB2 | Ovarian carcinoma JHU | TMT 10-plex (13) | Hu *et al.* (23)<br>(PDC000110) | MaxQuant executed<br>in-house |
| dB3 | AROM+ transgenic vs.<br>wild-type mice | LFQ | Vehmas *et al.* (24)<br>(PXD002025) | Vehmas *et al.* (24)<br>(PXD002025) |

#### Biological datasets

To demonstrate the practical application of PRONE on biologically meaningful data, we obtained one publicly available label-free and two TMT datasets. The first dataset, dB1, from a cell culture study of Li *et al.* (22) focuses on proteome dynamics during osteogenic differentiation of human periodontal ligament stem cells (hPDLSCs). The second, dB2, is a clinical dataset from the ovarian carcinoma Johns Hopkins University proteome study of Hu *et al.* (23). Lastly, dB3, an LFQ dataset of Vehmas *et al.* (24), explores the effects of high estrogen to androgen ratio on the mice liver proteome (**Table 1** and **Supplementary Methods**).

### Review and Selection of Normalization Methods for Proteomics Data

A selection of 17 normalization and two batch effect correction methods was made based on a comprehensive literature review (**Table 2** and **Supplementary Methods**). The work of Välikangas *et al.* (2), Callister *et al.* (4), Chawade *et al.* (1), Dubois *et al.* (6), and Kultima *et al.* (28), gave a systematic review of commonly used normalization methods in proteomics data. Notably, many of the methods applied on proteomics data have originally been developed for DNA microarray analyses. In contrast, the novel approaches RobNorm of Wang *et al.* (9) and two variants of Normics of Dressler *et al.* (10) were included as the approaches have only been assessed in their original publication and were specifically developed for proteomics data. Phil Wilmarth, a data analyst focusing on the analysis of TMT experiments, provided a detailed evaluation of three normalization methods GlobalMean, IRS, and TMM, available online (26). Furthermore, several other normalization methods were included from studies focused on normalizing proteomics data (7,8).

**Table 2:** Summary of the normalization methods and batch-effect correction methods. We summarized the selected approaches into five categories, based on the nomenclature proposed by Wang et al. (9). The column scale specifies if normalization was applied on raw or log2-transformed data, based on recommendations found in the literature for each respective normalization method. Of note, this is one of several parameters that can be easily configured in PRONE. More detailed descriptions of the normalization approaches and batch effect correction techniques are provided in the Supplementary Methods.

| Category | Approach | Abbreviation | Origin | Scale | Previous Evaluation Studies |
| --- | --- | --- | --- | --- | --- |
| Simple sample shifting | Mean / Median Normalization | - | Microarray | raw (7) | (1,2,6–10,28,29) |
|  | Global Intensity Normalization | GlobalMean, GlobalMedian | Microarray | raw (7,26) | (1,7,8,15,26) |
|  | Median Absolute Deviation Normalization | MAD | Proteomics | log2 (7) | (1,7) |
|  | Normics with Median Normalization | NormicsMedian | Proteomics | raw (10) | (10) |
| Sample-to-reference | Quantile Normalization | - | Microarray | log2 (7) | (1,2,4,6–8,10,28–30) |
|  | Linear Regression Normalization | Rlr, RlrMA, RlrMACyc | Microarray | log2 (7) | (1,2,4,7,8,28) |
|  | Local Regression Normalization | LoessF, LoessCyc | Microarray | log2 (7) | (2,4,7–10,28,29) |
|  | Trimmed Mean of M-Values Normalization | TMM | RNA-seq | raw (26,31) | (26) |
| Variance stabilizing | Variance Stabilizing Normalization | VSN | Microarray | raw (7,32) | (1,2,7–10,28,30) |
|  | Normics with VSN | NormicsVSN | Proteomics | raw (10) | (10) |
| Model-based | EigenMS | - | Metabolomics | log2 (33) | (2,5,9) |
|  | RobNorm | - | Proteomics | log2 (9) | (9) |
| Batch effect correction | Internal Reference Scaling | IRS | Proteomics | raw (26) | (15,26) |
|  | limma::removeBatchEffects | limBE | Microarray | log2 (32) | (32) |

### Evaluation of the normalization methods

#### Intragroup and -batch variation

Intragroup variation of all sample groups was measured using the pooled median absolute deviation (PMAD) for each normalization method since the PMAD is less affected by outliers than the intragroup pooled variance estimate or the pooled coefficient of variation (1). The PMAD of a sample group *g* is defined as the average median absolute deviation (MAD), calculated on the samples of group g, over all the proteins *i* = 1, …, n.

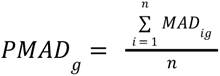

with *g_1_, …, g_e_* being the indices of the samples of condition *g*, and *e* being the number of samples of condition *g*. Additionally, the Pearson correlation coefficient was calculated for each pair of samples of a condition to determine the degree of similarity between the technical replicates in sample groups, with a high value indicating high intragroup similarity.

For the biological TMT datasets, diagnostic plots were used for visual evaluation, and a Silhouette coefficient-based alternative strategy to PMAD was implemented to quantitatively evaluate the performance of normalization and batch effect correction methods. The consistency of biological (condition) and technical (batch) sample groups was assessed for each normalization method using principal components (PCs) and the Silhouette coefficient, as described in (34). Initially, the Euclidean distance between all samples was computed based on the first three PCs, as in (35). Then, the Silhouette coefficient was used to quantify how well a sample fits within its assigned group (condition or batch) compared to other groups, with values close to 1 indicating correct group assignment and values close to -1 suggesting the sample belongs to a different group.

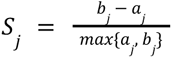

with *a*_*j*_ denoting the average Euclidean distance over the first three PCs of sample *j* and all other samples in the same group and *b*_*j*_ being the minimum average distance between sample *j* and samples in all other groups (34,35).

Finally, the average Silhouette coefficient for all samples within a (condition or batch) group was calculated to summarize the findings. In this context, a batch coefficient close to 1 indicates the presence of batch effects, while a condition coefficient close to 1 reflects a strong biological effect.

#### Differential expression analysis

DE of proteins was examined in each two-group comparison using limma (32) after the application of the different normalization methods in all spike-in and biological datasets. Furthermore, we applied the reproducibility-optimized test statistic (ROTS) (36) on spike-in datasets, as performed by Välikangas *et al.* (2), to evaluate the impact of the DE method on the assessment of normalization techniques. All p-values mentioned in this study have been adjusted per normalization method and dataset to control the false-discovery rate using the Benjamini-Hochberg procedure at a significance level of 0.05. In addition, a threshold of |logFC| > 1 was applied for the biological datasets.

Since the ground truth is known for the spike-in datasets, performance metrics such as area under the receiver operating characteristic curve (AUC), false positive rates (FPRs), and F1 scores were calculated. The calculation of an F1 score is particularly useful in cases with imbalanced datasets, such as in the spike-in datasets, as it integrates precision and recall into a single metric by using the harmonic mean. Since optimizing one metric can negatively influence the other, the F1 score provides a harmonized solution. In this context, a spike-in protein is counted as true positive when detected as DE while as false negative if not. In contrast, a background protein not DE, is counted as true negative, while one exhibiting significant DE is classified as false positive (FP).

Finally, since biological datasets lack a known ground truth, we compared the number of DE proteins and conducted an intersection analysis on the DE results to evaluate the consistency of DE proteins obtained with the different normalization methods. For this purpose, we calculated the Jaccard similarity coefficient of DE proteins between all pairs of normalization techniques, defined as the size of the intersection divided by the size of the union of the sets.

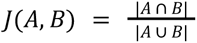

with A and B representing the sets of DE proteins identified by two normalization methods.

## Results

### PRONE: A competitive, accessible, and adaptable tool challenging state-of-the-art interfaces

#### Comparative evaluation of state-of-the-art tools

Open-source tools such as NormalyzerDE (7) and proteiNorm (8) were primarily designed to systematically assess normalization techniques. In contrast, proteoDA (37), tidyproteomics (29), AlphaPeptStats (30) offer an extensive range of functionalities for proteomics data analysis (**Table 3**). All five softwares are constrained in their data preprocessing features, such as missing value filtering and outlier detection. A collection of normalization techniques was curated from the literature (**Table 2**), yet none of the packages offer all the techniques (**Table 1**). Among the state-of-the-art tools, only AlphaPeptStats offers batch effect correction, restricting their analyses to datasets not measured in multiple batches. Although each tool integrates various DE methods, none allows the comparative assessment of normalization techniques on DE analysis. However, this is of high importance, since the choice of normalization technique has a direct influence on all downstream analyses (6,9,10). Moreover, the tools lack functionality for calculating performance metrics, such as FPRs and F1 scores, in spike-in datasets, thereby preventing the utilization of the known ground truth for evaluating normalization techniques. As most of the studies presenting a novel normalization method or assessing multiple normalization methods include at least one spike-in dataset for evaluation, a tool offering more functionalities specific for spike-in datasets is lacking.

**Table 3:**
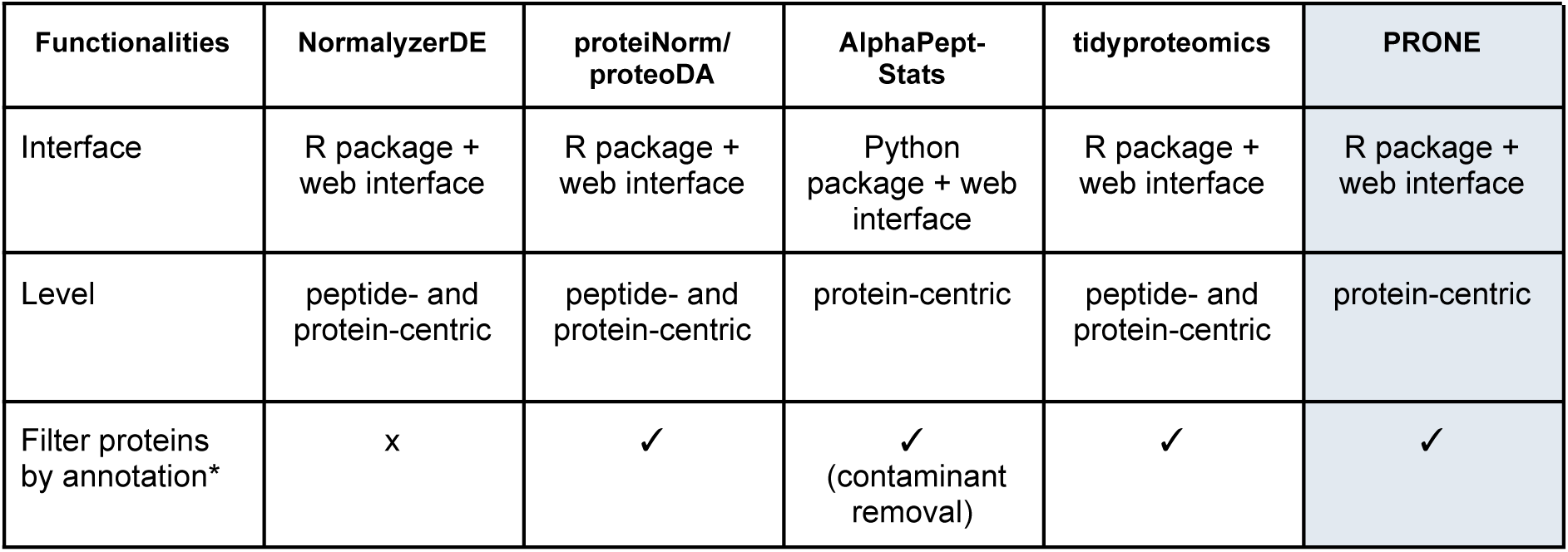
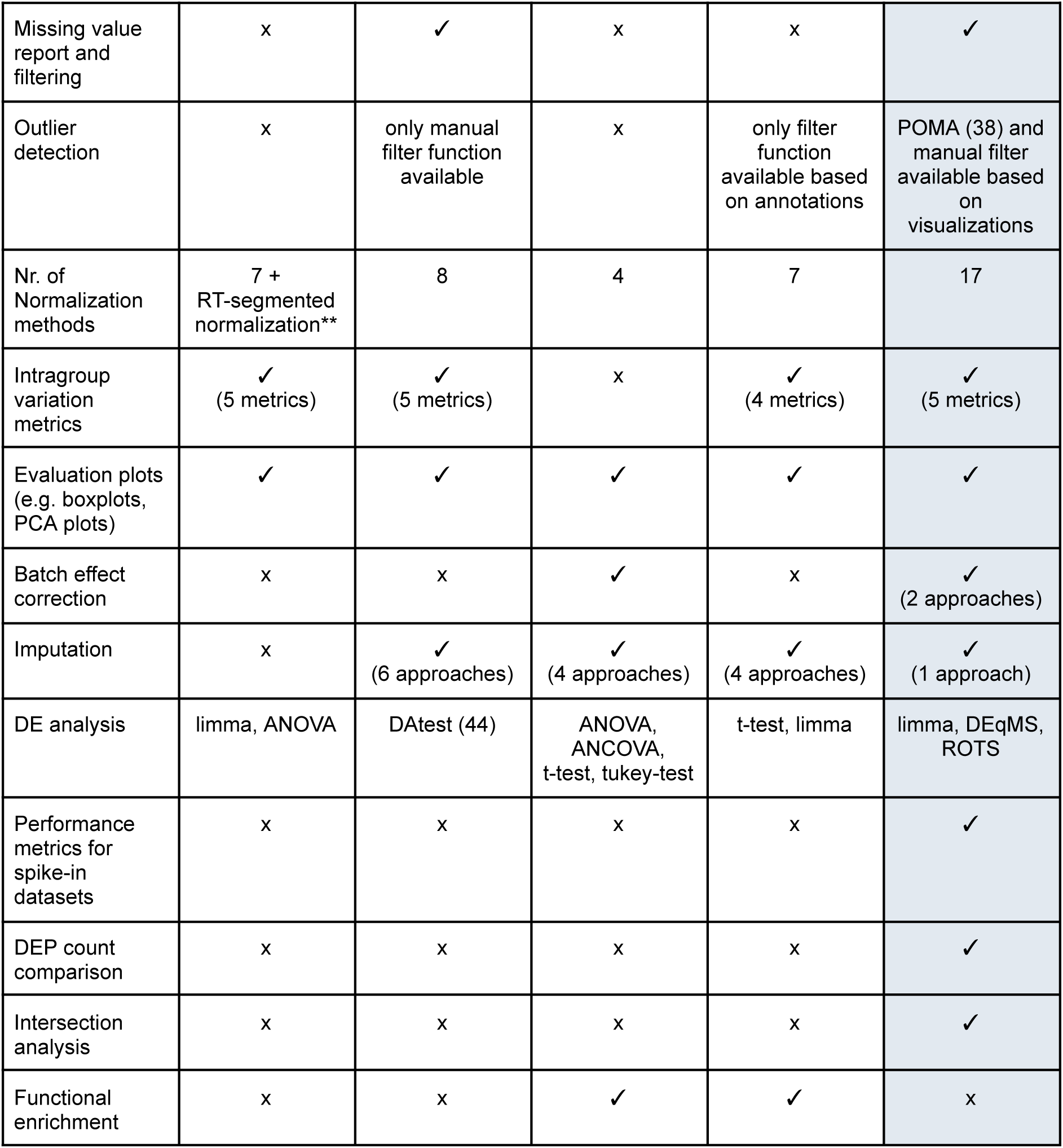
Comparative overview of publicly available state-of-the-art tools and PRONE. NormalyzerDE and proteiNorm, primarily designed for data normalization, and the tools proteoDA, AlphaPeptStats, and tidyproteomics, developed to facilitate proteomics data analysis, were compared along with PRONE (with light blue background) on various functionalities. Notably, since proteoDA is an extension of proteiNorm, these two were combined for comparison. *Filter proteins based on additional attributes, such as “Reverse” or “Potential contaminant”, as provided in MaxQuant output files. **In RT-segmented normalization, data points are grouped by their retention times into distinct segments, where normalization is performed individually within each segment to account for variations in electrospray ionization intensity (7).

| Functionalities | NormalyzerDE | proteiNorm/<br>proteoDA | AlphaPept-Stats | tidyproteomics | PRONE |
| --- | --- | --- | --- | --- | --- |
| Interface | R package +<br>web interface | R package +<br>web interface | Python<br>package + web<br>interface | R package +<br>web interface | R package +<br>web interface |
| Level | peptide- and<br>protein-centric | peptide- and<br>protein-centric | protein-centric | peptide- and<br>protein-centric | protein-centric |
| Filter proteins<br>by annotation* | x | ✓ | ✓<br>(contaminant<br>removal) | ✓ | ✓ |
| Missing value report and filtering | x | ✓ | x | x | ✓ |
| Outlier detection | x | only manual filter function available | x | only filter function available based on annotations | POMA (38) and manual filter available based on visualizations |
| Nr. of Normalization methods | 7 + RT-segmented normalization** | 8 | 4 | 7 | 17 |
| Intragroup variation metrics | ✓<br>(5 metrics) | ✓<br>(5 metrics) | x | ✓<br>(4 metrics) | ✓<br>(5 metrics) |
| Evaluation plots (e.g. boxplots, PCA plots) | ✓ | ✓ | ✓ | ✓ | ✓ |
| Batch effect correction | x | x | ✓ | x | ✓<br>(2 approaches) |
| Imputation | x | ✓<br>(6 approaches) | ✓<br>(4 approaches) | ✓<br>(4 approaches) | ✓<br>(1 approach) |
| DE analysis | limma, ANOVA | DAtest (44) | ANOVA, ANCOVA, t-test, tukey-test | t-test, limma | limma, DEqMS, ROTS |
| Performance metrics for spike-in datasets | x | x | x | x | ✓ |
| DEP count comparison | x | x | x | x | ✓ |
| Intersection analysis | x | x | x | x | ✓ |
| Functional enrichment | x | x | ✓ | ✓ | x |

#### Overview of PRONE

PRONE is a six-step workflow to benchmark the effectiveness of multiple normalization methods on protein-centric LFQ or TMT spike-in and biological proteomics data and guide the user’s decision on the normalization method (**Figure 2**). It offers the broadest selection of data normalization and batch effect correction techniques available.

**Figure 2:**
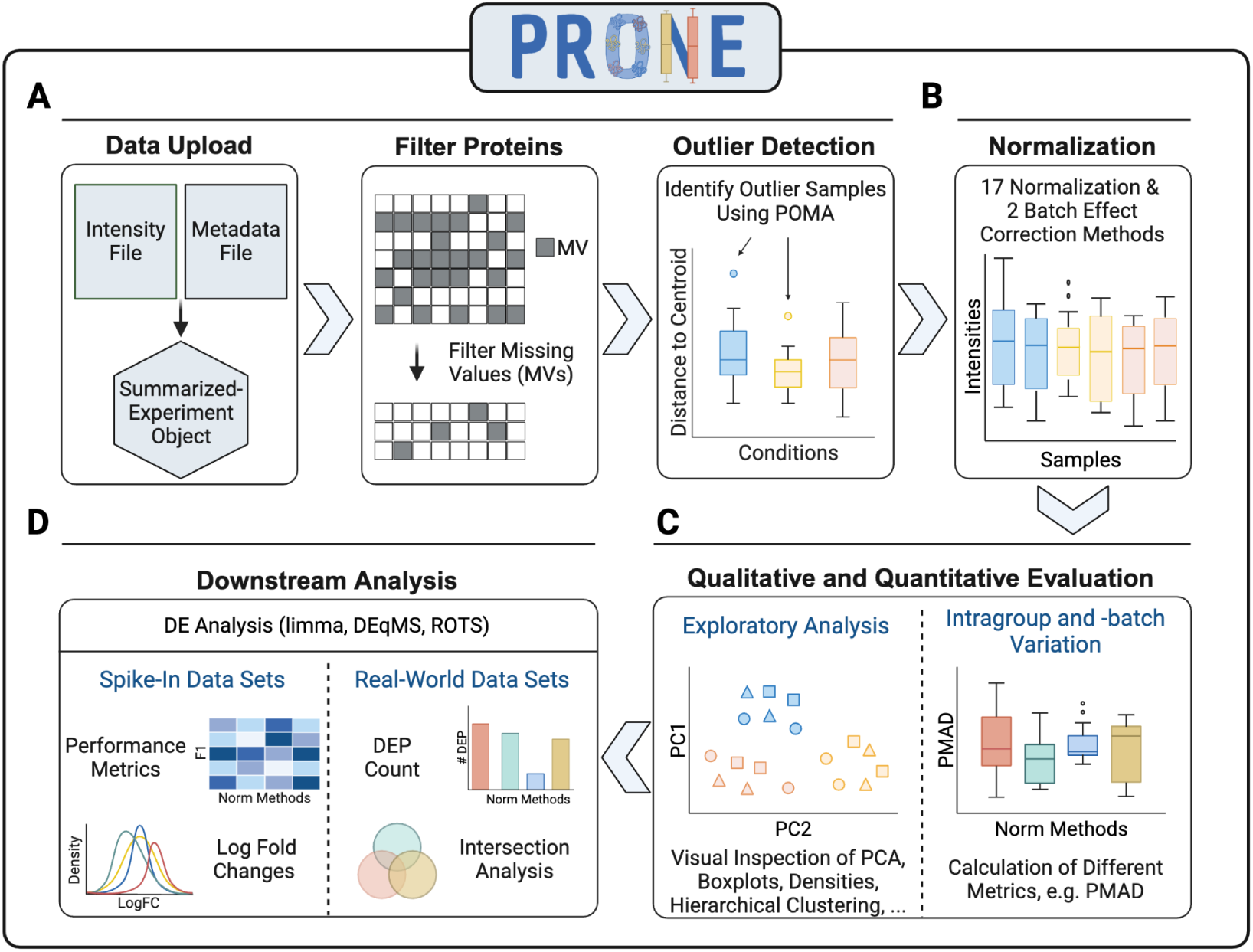
O**v**erview **of the six-step workflow of PRONE for proteomics data analysis and normalization method evaluation.** This six-step process initiates with the import of LFQ or TMT proteomic and two preprocessing steps (A). It allows for the exclusion of proteins exhibiting excessive missing values and the identification of outlier samples using POMA. The workflow includes 17 normalization methods and two batch effect techniques, which can be applied simultaneously and sequentially (B). Subsequent stages involve evaluating the performance qualitatively and quantitatively through the calculation of intragroup and -batch variation metrics and exploratory data analysis (C). Finally, three differential expression (DE) analysis methods are integrated to assess the efficacy of the applied strategies in detecting DE proteins (D). Thanks to the known ground truth of spike-in datasets, the evaluation of DE results is based on metrics, such as F1 scores. In contrast, the evaluation of DE results of biological datasets is based on intersection analyses of DE results obtained by the various normalization methods.

As input, PRONE requires two files: the raw protein intensity table and a metadata file (**Figure 2A**). PRONE is currently limited to protein-centric data, requiring the input protein intensity file to be in a tabular format, with proteins represented by rows and samples identified by columns. Other additional columns originating from MaxQuant (39), for instance, are allowed and can be accessed at any time.

The metadata file needs to specify, at minimum, the sample names and their corresponding group classifications.

Since the quality of MS-based proteomics data is challenged by the presence of missing values, users can specify a threshold for the minimal percentage of samples with a valid value for a given protein. Proteins not fulfilling this criterion can be filtered out. For researchers in favor of imputation rather than protein filtering, we provide a mixed imputation approach (40). This approach applies k-nearest neighbor imputation for proteins with missing values assumed to be missing at random, while values missing not at random are imputed using random draws from a left-shifted Gaussian distribution. A protein is classified as having missing values not at random if it exhibits missing values across all replicates of at least one condition. In addition, we included the multivariate outlier detection method of POMA (38). POMA computes the centroids of sample groups and employs a threshold on the interquartile range of each sample relative to its group centroid to identify outliers specific to each sample group. POMA outlier samples can be examined using various visualization techniques and subsequently either retained or excluded from the dataset.

PRONE provides considerable flexibility to users concerning normalization acknowledging the lack of universally accepted standards (**Figure 2B**, **Table 2**). For instance, batch effect correction can be executed either before or after normalization, and users decide whether to apply normalization on raw or log2-transformed data. The results of the normalization methods can then be compared using exploratory data analysis and visualizations, such as boxplots and PCA plots, or by calculating different intragroup variation metrics, e.g. pooled median absolute deviation (PMAD) (**Figure 2C**). Additionally, hierarchical clustering of samples and the computation of Silhouette coefficients based on PC components are provided to identify potential batch effects (see **Methods**) (14,34). Notably, PRONE offers the option to display evaluation metrics and data visualizations for a single normalization method at a time or to compare various normalization approaches in a single plot.

DE analysis (**Figure 2D**) can be conducted using limma (32), ROTS (36), or DEqMS (20). Given the known ground truth in spike-in datasets, various standard metrics, including F1 scores, can be employed to assess normalization techniques in detecting DE proteins. Furthermore, it is possible to compare the calculated logarithmic fold changes at base 2 (logFCs) against the expected logFCs derived from the known spike-in shifts. In biological data scenarios lacking a ground truth, alternative evaluation strategies are necessary. These include summarizing the counts of DE proteins across normalization methods, and conducting intersection analyses of DE findings. While DE intersection was previously conducted by other authors (10,28), it has not yet been incorporated into an R package or graphical user interface before PRONE. Jaccard similarities are calculated to evaluate the consistency of DE proteins identified using different normalization methods (see **Methods**).

Notably, PRONE provides features for pre-processing, normalization, evaluation of normalization techniques, and downstream analysis, which extends its utility beyond normalization as an integrated platform for the comprehensive analysis of proteomics data. PRONE is available as an R Bioconductor package and a user-friendly R Shiny App, accessible under https://exbio.wzw.tum.de/prone, to provide access to the functionalities for analyzing biological proteomics data without extensive programming knowledge.

### Spike-in datasets

We downloaded six publicly available spike-in datasets (**Table 1**) and used PRONE to compare the normalization methods in their ability to reduce intragroup variation and detect DE fold-changes (see **Methods and Supplementary Methods**). A standardized preprocessing strategy was applied to all spike-in datasets, encompassing protein filtering based on multiple criteria, such as missing value filtering and potential contaminant removal, and outlier sample detection (**Supplementary Table 1)**. Each dataset was subjected to all 17 normalization techniques (**Table 2**).

#### Impact of normalization on intragroup variation

Similar to previous evaluation studies (1,2,4,8,9), the PMAD and Pearson correlation coefficients were computed for each sample group within a dataset and across all normalization methods to assess their impact on intragroup variation and sample similarity. Normalization mainly decreased intragroup variation between technical replicates in all datasets compared to unnormalized log2-transformed data, except TMM normalization (**Figure 3A** and **Supplementary Table 2**). MAD normalization consistently achieved the highest percent reduction except for dataset dS2, for which EigenMS reduced intragroup variation the most (**Supplementary Table 2**). Similar results were obtained by the other normalization methods.

**Figure 3:**
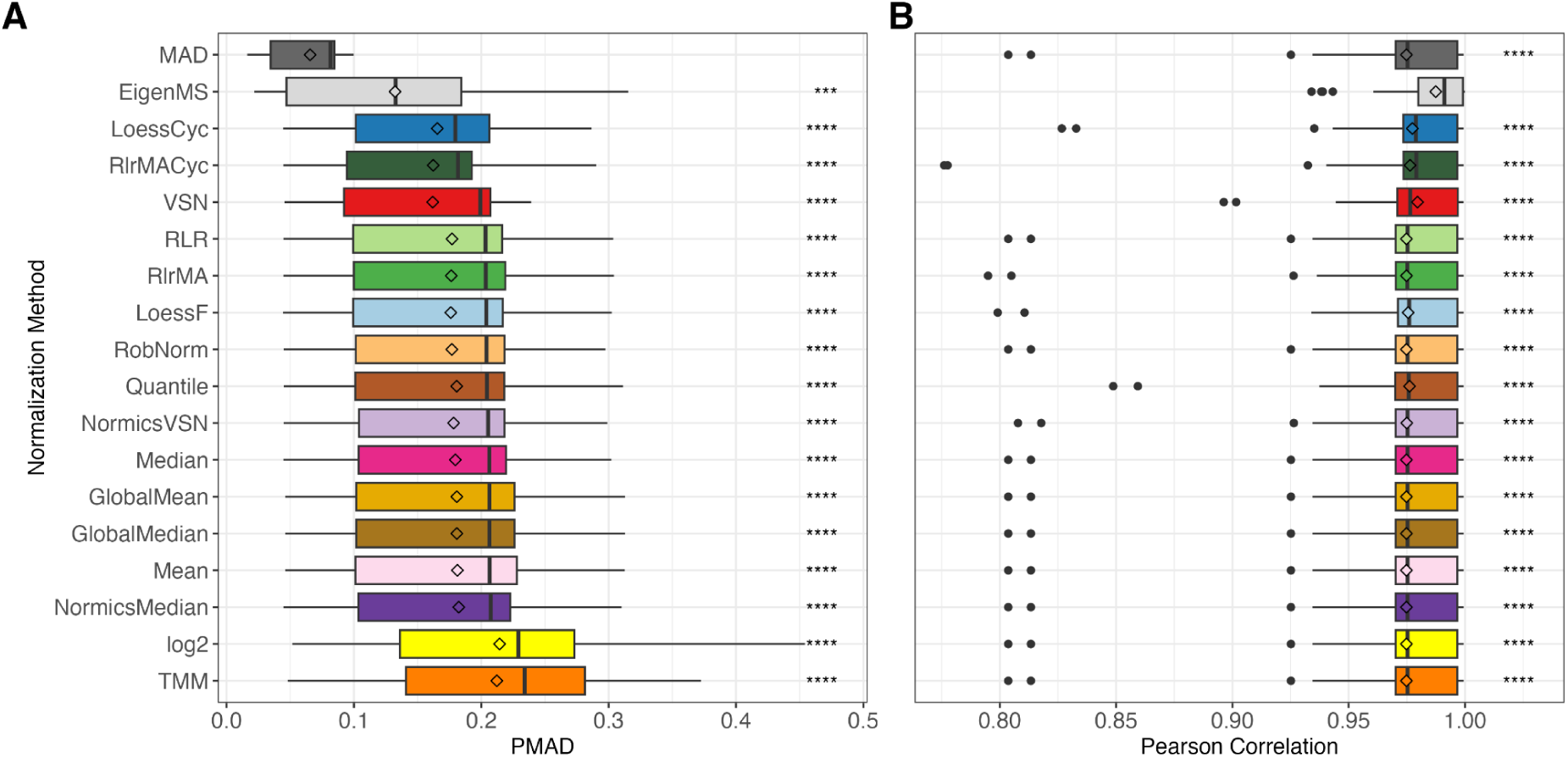
Overall intragroup variation and similarity of normalization methods in spike-in datasets. Boxplots depicting the distribution of pooled median absolute deviation (PMAD, A) and Pearson correlation coefficients (B) for all sample groups across all spike-in datasets, with normalization methods and unnormalized log2-transformation (‘log2’) ranked by their median PMAD. Diamonds represent mean PMAD and Pearson correlation in (A) and (B), respectively. Paired Wilcoxon rank sum tests were conducted to compare the PMAD and Pearson correlation values of MAD and EigenMS, respectively, with those of all other normalization techniques. Detailed results per spike-in dataset are provided in Supplementary Tables 2 and 3. *p ≤ 0.05, **p ≤ 0.01, ***p ≤ 0.001, **** p ≤ 0.0001.

Consistent findings were noticed across the spike-in datasets regarding the intragroup similarity between technical replicates measured using the Pearson correlation coefficient (**Figure 3B** and **Supplementary Table 3**). EigenMS was an exception and showed a most noticeable increase in correlation. In contrast to the observed differences in intragroup variation, MAD normalized intensities did not exhibit a substantial difference in sample correlation compared to other normalization methods. LoessCyc, VSN, and RlrMACyc had an overall higher median of Pearson correlation over the sample groups in all spike-in datasets.

#### Effect of normalization on differential expression

DE analysis was conducted for each spike-in dataset across all pairwise comparisons using limma (32) with multiple testing corrections using Benjamini-Hochberg procedure (41), setting an adjusted p-value threshold of 0.05. Since prior research predominantly assessed the DE results of normalization approaches using AUC values (2,7,9), we calculated the AUC for every pairwise comparison and each spike-in dataset (Figure 4A). The AUC analysis suggests high levels of performance, as indicated by median AUC values exceeding 0.75 across all normalization methods. However, in this study, we observed elevated FPRs in the spike-in datasets (Figure 4B), highlighting the limitations of relying solely on AUC values. Due to these observations and the imbalance of spike-in and background proteins (**Table 1**), we employed F1 scores for the conclusive assessment of the DE outcomes (Figure 4C **and 4D)**. The analysis revealed significant fluctuations in F1 scores across the different spike-in datasets (Figure 4D), e.g., the TMM and cyclic regression normalization methods demonstrated high performance in dataset dS6 but showed reduced efficacy in dataset dS2. Overall, RobNorm performed significantly better than all other methods (Figure 4C **and 4D**) except LoessF (Wilcoxon-test p-value = 0.1, **Supplementary** Figure 1). NormicsVSN and the linear regression methods RlrMA and RLR ranked among the top-performing methods based on the median F1 score across all spike-in datasets and pairwise comparisons (Figure 4C). In contrast, with EigenMS, the median F1 score is below 0.4, which aligns with its significantly higher FPRs (Figure 4B). Notably, in dS2, the linear and local regression methods employing a reference sample, including RlrMA, RLR, and LoessF, demonstrated overall high median F1 scores (> 0.6) compared to their cyclic counterparts (< 0.2) (Figure 4D). The poor performance of cyclic regression methods and EigenMS on dS2 can be attributed to the relatively high proportion of missing values in this dataset. Due to differing criteria for handling proteins with missing values (see **Supplementary Methods**), these methods can introduce additional missing values during normalization, which are subsequently classified as non-DE.

**Figure 4:**
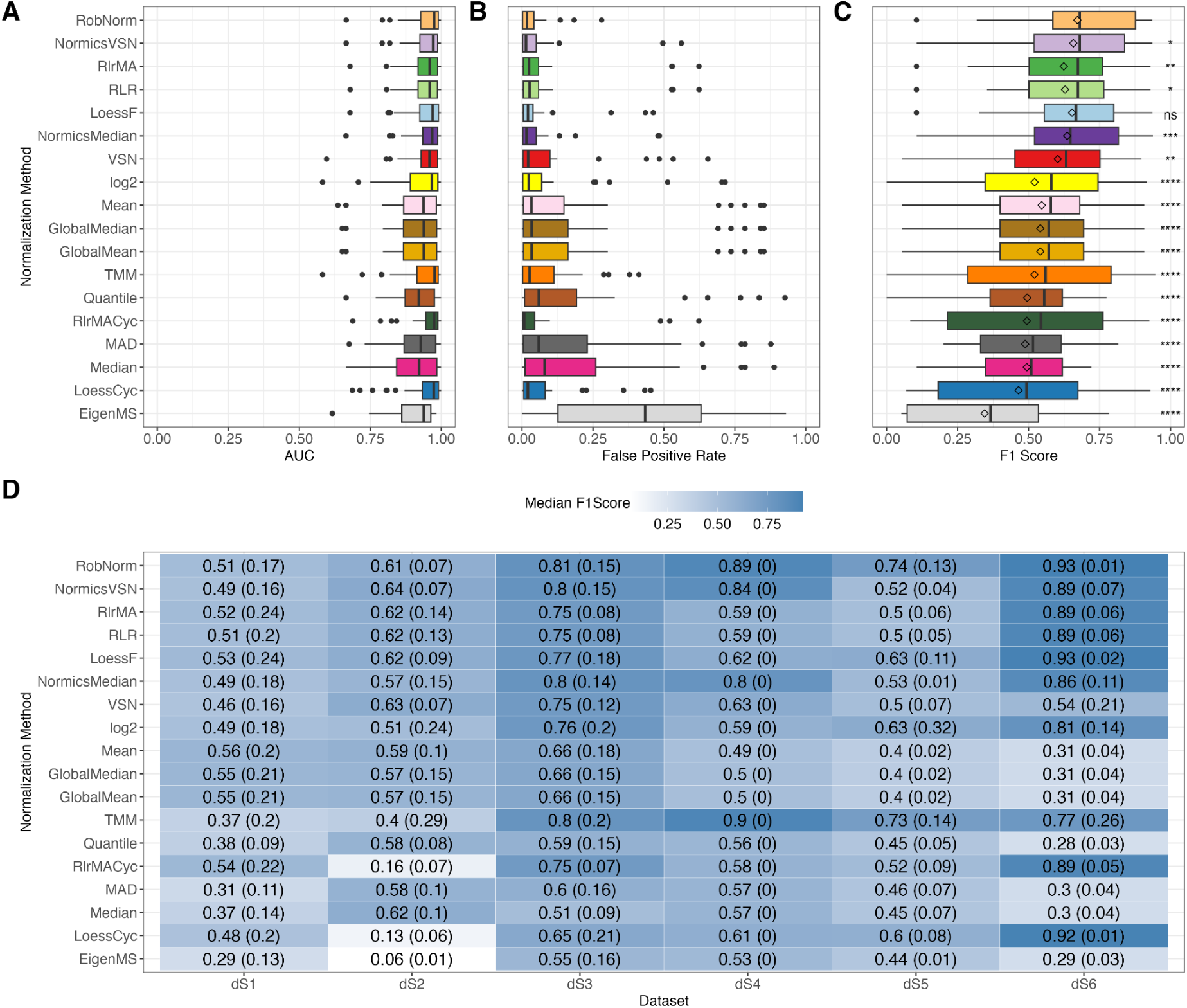
E**f**fect **of normalization on differential expression and comparison of evaluation metrics.** Differential expression analysis was conducted on every pairwise comparison of each spike-in dataset using limma with a threshold of 0.05 on Benjamini-Hochberg-adjusted p-values. Area under the receiver operating characteristic curve (AUC) values (A), FPRs (B), and F1 scores (C) were calculated for all normalization methods across all pairwise comparisons and spike-in datasets. Diamonds in panel C represent mean F1 scores. Paired Wilcoxon rank sum tests were conducted to compare the F1 scores of RobNorm with those of all other normalization techniques. Median F1 scores with the median absolute deviations indicated in brackets separated by spike-in dataset are shown in (D). The higher the F1 score, the better the performance of the normalization method. For all panels, normalization methods were sorted according to the median F1 score over all datasets. *p ≤ 0.05, **p ≤ 0.01, ***p ≤ 0.001, **** p ≤ 0.0001, ns = not significant.

#### Effect of normalization on log fold changes

Due to the experimental design of spike-in datasets, it is anticipated that the concentrations of the spike-in proteins will exhibit variability, whereas the concentrations of the background proteins are constant. The logFCs of the spike-in proteins were typically overestimated, both in the normalized data and in log2-transformed data (**Supplementary** Figure 2A). In general, the tested methods yielded comparable results in estimating logFCs of the spike-in proteins, with the exception of MAD normalization, which consistently produced higher logFCs of the spike-in proteins than expected. While the logFC values of the background proteins should remain unchanged since their concentrations remain constant across the sample groups, the logFC distributions of the background proteins across all datasets were not centered at zero for the majority of the normalization approaches (**Supplementary** Figure 2B). Nonetheless, the distributions of the logFCs in datasets normalized using RobNorm, LoessF, and LoessCyc, were notably concentrated around zero. Of these, RobNorm demonstrated the most accurate outcome (median logFC = 0.00109).

#### Impact of differential expression method on proteomics data analysis

To address the observation that limma tends to detect a high amount of FPs in spike-in datasets (Figure 4B), we integrated another method, reproducibility-optimized test statistic (ROTS) test, into the pipeline. This integration was based on prior studies demonstrating ROTS’ enhanced performance in detecting fold-differences over the conventional t-test in proteomics data analysis (2,36,42). While PRONE also includes DEqMS as an alternative to limma and ROTS, its performance was not evaluated in this study due to the limited availability of peptide counts per protein in the majority of the spike-in datasets.

Overall, ROTS consistently reduced the number of FPs across all normalization techniques (**Supplementary** Figure 3A). The reduction in FPs by ROTS was associated with a slight decrease in the number of TPs in various normalization methods. These variations in the counts of FPs and TPs influenced the ranking of normalization methods when evaluated by median F1 scores across all pairwise comparisons and spike-in datasets (**Supplementary** Figure 3B). The cyclic regression methods demonstrated improved performance following the application of ROTS, and LoessF emerged as the best-performing method based on the median F1 score followed by RobNorm, NormicsVSN, NormicsMedian, and VSN, though the differences in performance were not significant (Wilcoxon-tests p-values > 0.05). Despite these alterations, the methodologies that previously showed best performance according to F1 scores based on limma DE results, specifically RobNorm and NormicsVSN, continued to rank among the top-performing methods with the adoption of ROTS. Finally, the F1 scores of ROTS and limma were compared in a paired Wilcoxon rank sum test per normalization technique (**Supplementary** Figure 3C). For certain normalization methods, i.e., TMM, RlrMACyc, and LoessCyc, significant differences were observed in the F1 scores between ROTS and limma, whereas for the best-performing methods, LoessF, RobNorm, and NormicsVSN, no significant differences were detected. Thus, while the choice of DE method influences the performance evaluation of the normalization approaches, it does not substantially alter the ranking of the best performing methods in our specific cases.

### Biological datasets

While spike-in datasets offer a known ground truth for method validation, the absence of such ground truth in biological datasets complicates the selection of an appropriate normalization technique. To illustrate the application of PRONE in typical biological study settings, we selected one LFQ and two TMT datasets (see **Methods** and **Table 1**). Following the preprocessing of the datasets with PRONE (**Supplementary Table 4**), we primarily focused on evaluating intragroup variation and comparing DE results. Based on the recommendations of (10,43), we evaluated VSN and NormicsVSN with the default lts.quantile value of 0.9, as well as with the lts.quantile parameter set to 0.5, hereafter referred to as VSN_0.5 and NormicsVSN_0.5 in this study.

#### Impact of normalization on intragroup and -batch variation

In the cell culture dataset dB1 (**Table 1**), we examined the biological replicates across the four time points. Initially, we normalized the dataset using PRONE’s 17 normalization approaches. Hierarchical clustering of samples revealed clear evidence of batch effects according to TMT-labeling for all normalization methods, except for EigenMS (**Supplementary** Figure 4A). The application of batch effect correction methods, IRS and limBE, on normalized data mitigated this technical variability to some extent. Notably, the biological signal, corresponding to timepoints, became more pronounced with limBE for the majority of normalization techniques (**Supplementary** Figure 4B). However, since diagnostic plots rely on visual interpretation, quantitative approaches offer a more objective means of evaluating whether batch effects have been effectively removed while preserving the underlying biological signal. Given the limitations of the PMAD metric in the spike-in datasets, an alternative approach based on PCs and Silhouette coefficients was used (see **Methods**). Silhouette coefficients are a measure for the goodness of clustering, for which a high value denotes that data points belong to the tested sample group, e.g. batch or condition. A batch coefficient close to 1 suggests the presence of batch effects, whereas a condition coefficient close to 1 indicates a strong biological effect. The Silhouette coefficients of TMT-batch groups were significantly reduced following the application of either IRS or limBE, a trend observed across all normalization methods except EigenMS (**Supplementary** Figure 5). For simplicity, we summarized the Silhouette coefficients for all normalization methods with limBE showing the most substantial reduction in median Silhouette coefficient overall (Figure 5A). Notably, similar to the results observed using the hierarchical clustering, the biological signal, e.g., related to different time points, remained preserved in the data after correcting for batch effects (Figure 5B).

**Figure 5:**
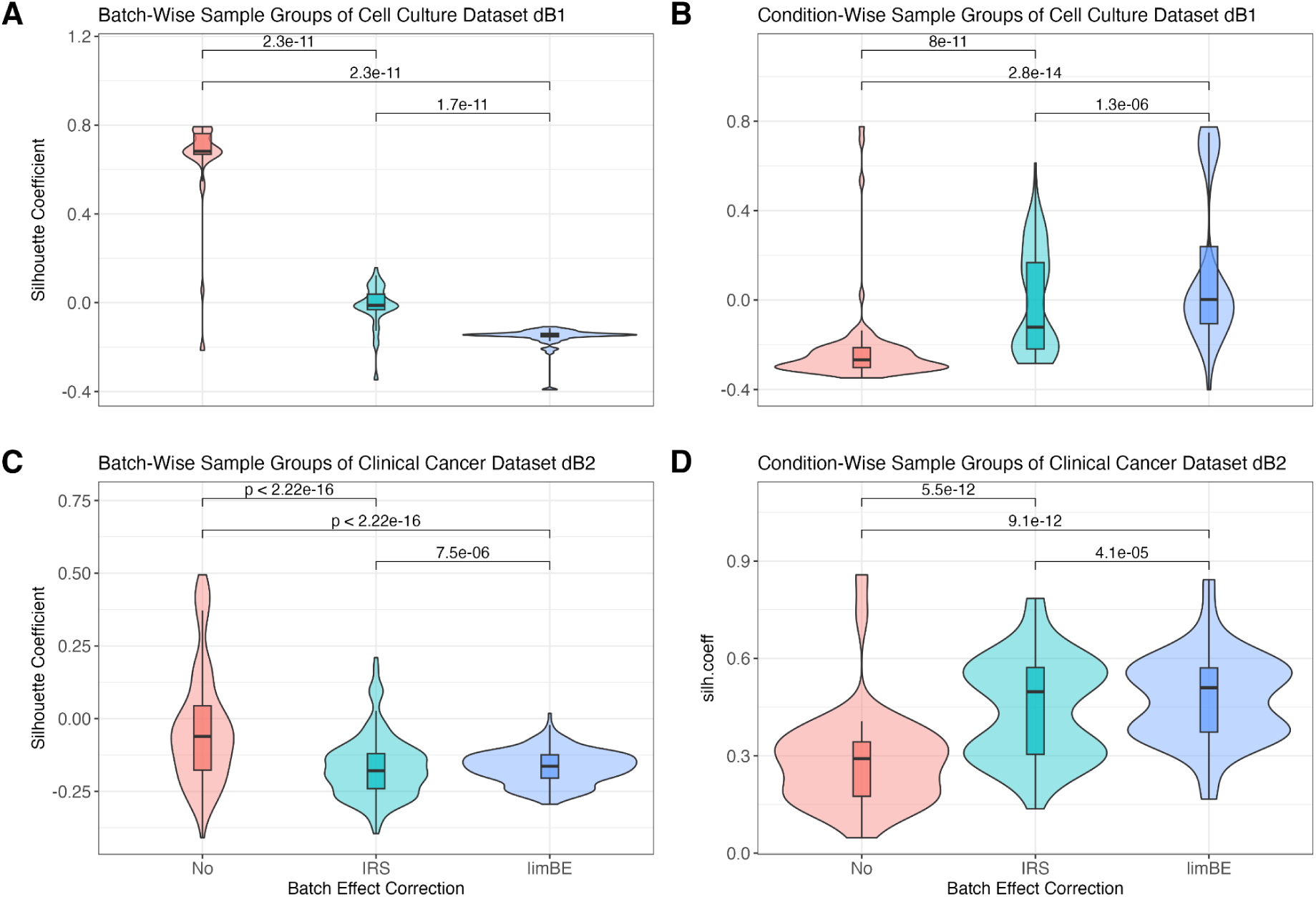
B**a**tch **effect assessment of the biological TMT datasets using Silhouette coefficients per normalization method.** The Silhouette coefficients, based on the Euclidean distances among samples derived from the first three principal components, quantify how well each sample fits within its assigned group. The Silhouette coefficients were computed for each sample group, batch- and condition-wise across all combinations of normalization techniques and batch effect correction techniques in the cell culture dataset dB1 (A, B) and the clinical cancer dataset dB2 (C, D), respectively. Normalization methods were applied without batch effect correction (No), with internal reference scaling (IRS), and with limma::removeBatchEffects (limBE). Paired Wilcoxon rank sum tests were conducted to compare the Silhouette coefficients of the batch effect correction methods.

This finding was further validated using the clinical cancer dataset dB2. Due to the comparatively large sample size and heterogeneity of cancer samples, we applied hierarchical clustering of samples based on protein expression levels, rather than relying on the visual evaluation of PCA plots, in conjunction with quantitative evaluation using Silhouette coefficients. The hierarchical clustering of samples showed that without batch effect correction, samples exclusively group by batch, while batch effect correction mitigates this effect (**Supplementary** Figure 6). Notably, samples normalized with EigenMS cluster by pathological status rather than batch effect without the use of any batch effect correction method. Furthermore, as already observed in the cell culture dataset dB1, limBE efficiently removes the batch effects and makes the biological signal more prominent, which is in concordance with the Silhouette coefficients (Figure 5C, Figure 5D **and Supplementary** Figure 7).

For completeness, the intragroup variation metric PMAD used in the spike-in datasets, was computed on the biological TMT datasets as well (**Supplementary** Figure 8). The results align with the Silhouette coefficients (Figure 5). Consequently, we proceeded with the downstream analysis using data from all normalization methods following the application of limBE. In contrast, for the LFQ dataset dB3, all normalization methods were applied without any batch effect correction as samples were not measured in different batches.

#### Impact of normalization on DE results of biological proteomics data

For the sake of simplicity in analyzing dB1 we, next, focused on the pairwise comparison for the longest experimental time span of the cell culture experiment (D0-D14). Note, that all other pairwise comparisons of dB1 are included in the supplement (**Supplementary** Figure 9). We calculated the number of up- and down-regulated DE proteins using an absolute logFC threshold of 1, and normalization by TMM, EigenMS, MAD, and VSN showed high variability in the number of DE proteins compared to the other methods (Figure 6A). Additionally, we observed that applying batch effect correction on log2-transformed data (denoted as “log2”) yielded results comparable to those obtained using EigenMS and TMM normalization. EigenMS, TMM, and log2 resulted in over 125 DE proteins, whereas less than 25 DE proteins were detected by applying MAD or VSN. Interestingly, the number of DE proteins identified by normalization with VSN_0.5 but not default VSN was similar to the most normalization techniques. The results were further examined by generating volcano plots using PRONE, which revealed that the distribution of p-values and logFC values from log2 and TMM normalization are of distinct parabolic shape that are highly pronounced compared to the distribution from the other normalization methods. Furthermore, the results from log2, TMM, and EigenMS normalization were skewed towards significantly higher logFC values compared to the other methods (**Supplementary** Figure 10).

**Figure 6:**
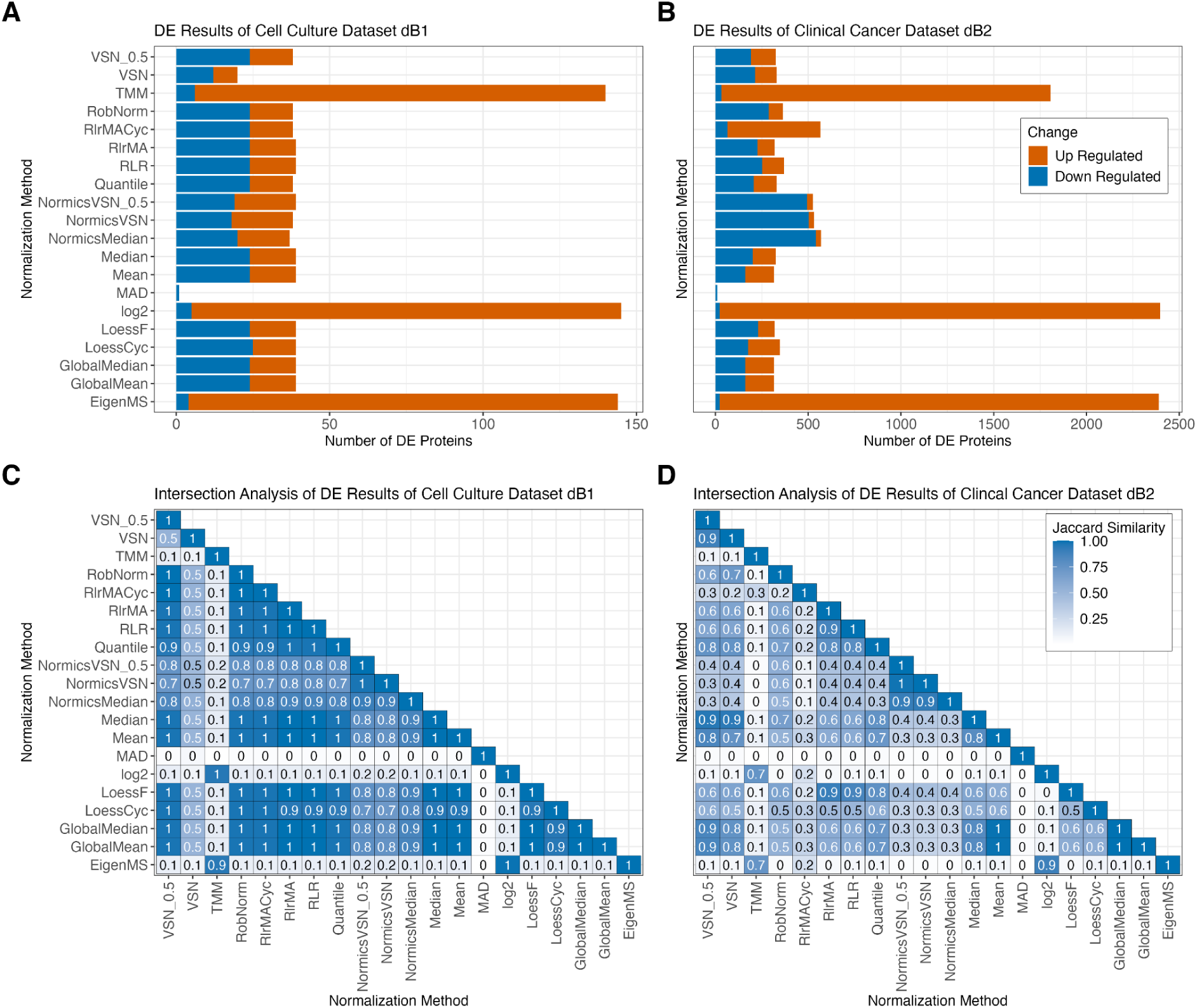
D**i**fferential **expression results of biological TMT datasets.** Bar plots showing the number of differentially expressed (DE) proteins of the pairwise comparison D0-D14 for each normalization method with limBE applied on top to remove batch effects in the cell culture dataset dB1 (A) and the clinical cancer dataset dB2 (B), respectively. DE proteins are classified as up- and down-regulated using |logFC| > 1 and Benjamini-Hochberg adjusted p-value < 0.05 was applied, revealing that the normalization method affects downstream analysis results. Heatmaps displaying Jaccard similarity coefficients for DE gene sets identified by all pairs of normalization methods for the cell culture dataset dB1 (C) and for the clinical cancer dataset dB2 (D), respectively. The values within the tiles were rounded to one decimal place.

The clinical cancer dataset dB2 revealed higher variations across all normalization techniques (Figure 6B). However, similar to the cell culture dataset dB1, log2, normalization by TMM and EigenMS resulted in significantly more DE proteins, predominantly up-regulated, whereas the MAD method detected almost no DE proteins. Notably, normalization by NormicsMedian and NormicsVSN showed a higher proportion of down-regulated DE proteins compared to other normalization techniques. In the LFQ dataset dB3, normalization by EigenMS and MAD resulted in the highest and lowest number of identified DE proteins, respectively (**Supplementary** Figure 11A).

While the amount of identified DE proteins is similar for the majority of normalization techniques for the cell culture dataset dB1 and LFQ dataset dB3, minor variations could potentially influence downstream analyses such as network and gene set enrichment. PRONE introduces a novel feature that assesses the DE protein sets: intersection analyses of the DE results based on Jaccard similarity coefficients. Apart from the methods with high variability in the number of DE proteins in the cell culture dataset dB1 such log2, VSN, TMM, MAD, and EigenMS, the same proteins were identified as DE independent of the normalization method (Jaccard coefficients above 0.9) (Figure 6C and **Supplementary** Figure 12). The exceptions are NormicsMedian and the two NormicsVSN approaches, which exhibited comparatively lower coefficients. The clinical cancer dataset dB2 revealed much lower Jaccard similarity coefficients across all normalization techniques, a contrast not as evident in earlier evaluation stages (Figure 6D). And the LFQ dataset dB3 showed results comparable to the cell culture dataset, with Jaccard coefficients exceeding 0.7 for all normalization methods except MAD (**Supplementary** Figure 11B).

However, we observed that MAD normalization generates notably lower absolute logFC values (**Supplementary** Figure 10 and 13A**-C**), explaining the reduced number of DE proteins identified under the applied logFC threshold for all datasets. To account for this, we applied a reduced logFC threshold of 0.5 to MAD, which not only increased the number of DE proteins but also improved the Jaccard coefficients, indicating greater overlap of DE proteins with those obtained using other normalization techniques (**Supplementary** Figure 13D-G).

## Discussion

### Evaluation of normalization methods

All normalization techniques, with the exception of TMM, reduced intragroup variation compared to unnormalized data, with EigenMS and MAD being most effective in this regard. MAD was not extensively evaluated before but the findings of EigenMS were confirmed in (2,9). Besides MAD and EigenMS, the linear, local regression methods and VSN rank among the most effective techniques for reducing intragroup variation, as demonstrated in (1,2,28). Additionally, with biological data, we demonstrated that batch effect correction is essential for TMT datasets to decrease the technical effect while maintaining the biological signal, as previously confirmed in (14,26,44) (Figure 7). The analysis of the two biological TMT datasets, dB1 and dB2, demonstrated that using the reference samples with IRS for batch effect correction was less effective compared to the model-based approach of limma.

**Figure 7:**
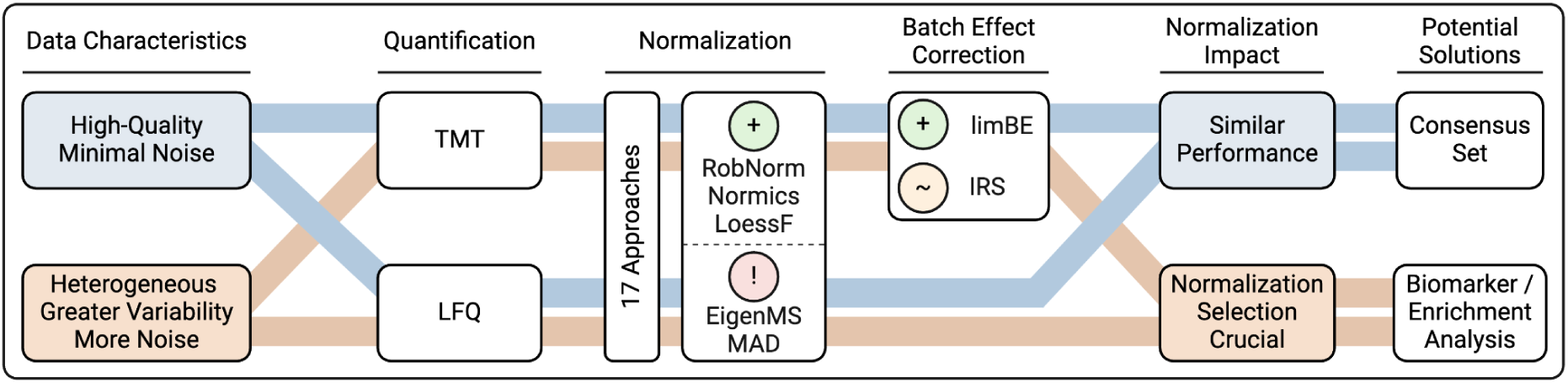
S**u**mmary **of evaluation study findings.** Our evaluation demonstrated that the impact of normalization on the proteomics data analysis is highly dependent on the underlying data characteristics, such as differences between high-quality datasets with minimal noise and heterogeneous datasets exhibiting greater variability. Among the evaluated methods, the normalization methods RobNorm, Normics, and LoessF showed most promising results on the presented spike-in and biological datasets, with limBE being particularly effective in reducing batch effects present in TMT datasets. Methods were classified as follows: a green plus for most accurate performance, a red exclamation mark for notable limitations, and an orange tilde for methods being less effective than those with a green plus classification.

Next, we evaluated the impact of normalization techniques on DE analysis. With spike-in data and biological data, we showed that compared to state-of-the-art practice, a reduction of intragroup variation is not directly related to the effectiveness of the normalization methods according to the DE results. Previous evaluation studies that did not exclusively focus on intragroup variation metrics but also employed DE analysis, partially confirmed these findings (2,9). Furthermore, we demonstrated that EigenMS-normalized data resulted in a high number of FPs in spike-in data. Generally, as already shown in (10), the identification of a high proportion of FPs is observed, which is not recognizable when using AUC values due to the class imbalance. Thus, in contrast to previous studies (2,7,9), we employed F1 score which uses precision instead of specificity (as in AUC) and thus gives more attention to the problem of false positives in detecting DE proteins. According to the median F1 score calculated over all pairwise comparisons and spike-in datasets, RobNorm is the best-performing normalization method, closely followed by LoessF which showed no significant difference in performance. Furthermore, NormicsVSN, which has not been previously independently evaluated, ranked among the top-performing methods. In contrast, EigenMS produced the lowest F1 scores, with a comparatively high number of FPs. Finally, when employing another test statistic for DE analysis, ROTS, the number of FPs decreased, with only a minimal impact on the number of TPs. Notably, certain methods exhibited a greater benefit from the use of ROTS compared to others, resulting in minor alterations in the ranking of the normalization methods. Although LoessF ranked highest with ROTS, its F1 scores were not significantly higher than those resulting from RobNorm, the Normics approaches, and VSN.

Since there is no ground truth for the three biological datasets, we based our evaluation on the number of DE proteins. While data normalized using TMM and EigenMS resulted in a substantially higher number of DE candidates, MAD resulted in a noticeably lower number of DE results. Based on the previous results on spike-in datasets, we hypothesize that the majority of DE proteins stemming from EigenMS may be FPs. The compositional correction achieved through TMM normalization was found to produce comparable results to those obtained without applying explicit normalization before batch effect correction, i.e., on log-transformed data (log2). While data normalized with MAD resulted in relatively low F1 scores in the spike-in datasets, the application of a less stringent logFC threshold in the DE analysis of biological datasets revealed a high overlap in identified DE proteins across other normalization methods (**Supplementary** Figure 13). However, the fact that MAD normalization aligns the data to a common scale with reduced variance leading consistently to lower intragroup variation and lower logFCs, may represent a potential limitation of the method. Hence, the evaluation of the presented spike-in and biological datasets highlighted certain limitations of the EigenMS and MAD approaches, while showing the importance of exploring and assessing emerging methods like RobNorm and the Normics approaches that were specifically developed for application in proteomics research (Figure 7).

Intersection analysis of the DE proteins in the cell culture dataset dB1 revealed that most normalization methods consistently produced comparable results. Conversely, the results of the clinical cancer dataset dB2 demonstrated that normalization has a substantial impact on downstream analyses and, hence on the overall conclusions drawn from proteomics data analysis. Hence, based on these two datasets, we hypothesize that for high-quality datasets with minimal noise, most normalization approaches perform similarly, while for more heterogeneous datasets with greater variability and noise, the choice of normalization method significantly affects the results (Figure 7). These findings, combined with the F1 scores of the spike-in datasets, indicate that the impact of normalization methods is highly dataset-specific.

Given the dataset-specific effects and the unknown ground truth in biological datasets, it is advisable to conduct a comparative evaluation of multiple normalization techniques, incorporating both intragroup variation metrics and DE analysis, to guide decisions toward appropriate normalization techniques. However, domain-specific knowledge will be indispensable to account for the dataset specificity depending on the biological questions.

### Study limitations and future work

Despite the comprehensive range of normalization methods and datasets, this study is subject to potential limitations. We focused our benchmark primarily on label-free and TMT datasets despite the widespread use of other quantification strategies in the proteomics community, e.g., SILAC, iTRAQ, ICAT, and SILAM (45,46). However, given the minimal data requirements of PRONE, other types of proteomics datasets could be readily incorporated into future evaluations. Additionally, the study was restricted to TMT spike-in datasets measured within a single batch, since no suitable publicly available multi-batch spike-in TMT datasets could be found during data collection. Moreover, we focused on protein-centric normalization although proteomic data can be normalized also on peptide levels, as benchmarked in (4,28). In this study, we chose to apply POMA for outlier detection prior to normalization, as outliers can substantially influence the normalization process. Nonetheless, a thorough evaluation of the interplay between sample outlier detection and normalization would be of considerable value for future research. Likewise, the systematic evaluation of imputation methods in combination with normalization techniques on DE results provides another reasonable future extension of PRONE. Although this study compares the DE methods limma and ROTS, PRONE also incorporates DEqMS, a DE method developed by Zhu *et al.* (20) that reduces variance based on the number of peptides used for quantitation. The performance of DEqMS was not investigated in this study, as the peptide-count per protein was only available for a minority of the spike-in datasets. However, since DEqMS is already implemented in PRONE, it can be applied and compared to the other DE methods. Another option to extend future versions of PRONE beyond DE analysis is to offer functional enrichment techniques and the evaluation of the impact of the normalization methods on enrichment results, as performed in (9,10). This could be particularly beneficial for heterogeneous datasets with higher variability, such as the clinical cancer dataset dB2, where normalization has been shown to exert a greater impact on the DE results. Incorporating functional enrichment would provide biological interpretation on the findings (Figure 7). In contrast, for high-quality datasets with minimal noise, such as the cell culture dataset dB1, we hypothesize that utilizing a consensus set of DE proteins could serve as a viable approach for downstream analyses. However, we emphasize that this hypothesis requires further consideration and validation.

Despite some limitations in this study, the PRONE package already incorporates many key functionalities necessary for further analyses and result evaluation. Importantly, this study represents an important step toward establishing a universal pipeline for proteomics data analysis, with a particular emphasis on normalization.

## Data Availability Statement

The proteomics data of the spike-in and biological datasets without known ground truth utilized in this study and in the vignettes of PRONE are available through Zenodo at 10.5281/zenodo.12657423.

## Code Availability Statement

The R package and Shiny app source code can be accessed on GitHub at https://github.com/daisybio/PRONE and https://github.com/daisybio/PRONE.Shiny, respectively. Both are distributed under the GPL-3.0 license. Furthermore, for those interested in the source code of the evaluation study, the corresponding R code is available at https://github.com/daisybio/PRONE.Evaluation/.

## Funding

This work was supported by the Deutsche Forschungsgemeinschaft (DFG, 516188180), the German Federal Ministry of Education and Research (BMBF) within the framework of the *e:Med* research and funding concept (*grants 01ZX1910A, 01ZX1910D, 01ZX2210A and 01ZX2210D*), “CLINSPECT-M-2” (grants 161L0214A and 16LW0243K) and preclinical confirmatory study framework (REGAGforBone, grant: 01KC2304C). T.L. was awarded with a seed funding under the Excellence Strategy of the Federal Government and the Länder. Contributions by J.K.P. were partly funded by the Bavarian State Ministry of Education and the Arts in the framework of the Bavarian Research Institute for Digital Transformation (bidt, grant LipiTUM).

**Lis Arend** is a PhD candidate at the Technical University of Munich in the group of Prof. Dr. Markus List.

**Klaudia Adamowicz** is a PhD candidate at the University of Hamburg in the group of Prof. Dr. Jan Baumbach.

**Johannes R. Schmidt** is a post-doctoral researcher at the Fraunhofer Institute for Cell Therapy and Immunology (IZI) in the group of Prof. Dr. Stefan Kalkhof. He obtained his PhD at the Leipzig University and Helmholtz Centre for Environmental Research (UFZ) in Leipzig.

**Yuliya Burankova** is a PhD candidate at the Technical University of Munich (TUM), supervised by Prof. Dr. Bernhard Kuster (TUM) and Prof. Dr. Jan Baumbach (University of Hamburg).

**Olga Zolotareva** leads the Computational and Systems Medicine group at the Institute for Computational Systems Biology led by Prof. Dr. Jan Baumbach at the University of Hamburg.

**Olga Tsoy** leads the Computational Genomics and Transcriptomics group at the Institute for Computational Systems Biology led by Prof. Dr. Jan Baumbach at the University of Hamburg.

**Josch K. Pauling** has been leading the research group LipiTUM at the Technical University of Munich and he is the group leader of the Computational Integrative Omics in Biomedicine group at the Institute of Clinical Chemistry and Laboratory Medicine (IKL) at the University Hospital and Faculty of Medicine Carl Gustav Carus of the Dresden University of Technology.

**Stefan Kalkhof** has been appointed as a professor for instrumental bioanalysis and serves as the director of the Institute for Bioanalysis at the University of Applied Sciences Coburg. He is also the head of the Proteomics Unit at the Fraunhofer Institute for Cell Therapy and Immunology (IZI).

**Jan Baumbach** is a full professor and director of the Institute for the Computational Systems Biology (CoSy.Bio) at the University of Hamburg. He obtained his PhD at the Center for Biotechnology in Bielefeld and was postdoctoral fellow at the University of California at Berkeley.

**Markus List** is an assistant professor of Data Science in Systems Biology at the Technical University of Munich. He obtained his PhD at the University of Southern Denmark and was a postdoctoral fellow at the Max Planck Institute for Informatics in Saarbrücken.

**Tanja Laske** is a group leader at the Institute for Computational Systems Biology led by Prof. Dr. Jan Baumbach at the University of Hamburg. She obtained her PhD at the Otto von Guericke University and Max Planck Institute in Magdeburg.

## Acknowledgments

The authors thank Alexander Dietrich, Quirin Manz, and Nicolas Trummer for their assistance in the deployment of PRONE. Additionally, we acknowledge Tommy Välikangas for supplying the data utilized in their evaluation study and Cecilia Jensen for her aid in sourcing suitable biological datasets. Figure 1 and 2 were created with BioRender.com.

## Author Contributions

L.A. and T.L. conceived and designed this study. J.K.P., S.K., J.B., M.L., and T.L. obtained funding and supervised the project. L.A. performed the bioinformatics analyses and visualizations with input from all authors. L.A. implemented the R package and R Shiny App PRONE with guidance from K.A. K.A. implemented the Normics normalization approach in R. J.R.S. and Y.B. assisted in re-analyzing raw MS data using MaxQuant. All authors provided critical feedback and discussion and assisted in interpreting the results, writing the manuscript, and improving the web service.

## Keypoints

● A total of 17 normalization techniques and two batch effect correction methods were systematically assessed in this study across six spike-in and three biological proteomics datasets without known ground truth.
● RobNorm and both variants of Normics, which have not been previously evaluated independently, performed consistently well based on F1 scores across spike-in datasets.
● Differential expression analysis of biological datasets without known ground truth revealed that the choice of normalization has an impact on downstream analyses and the impact is dataset-specific.
● We introduced PRONE, the PRoteomics Normalization Evaluator, an R Bioconductor package coming along with a graphical interface to streamline a proteomics data analysis, especially focusing on data normalization.

