## Supplementary for "Systematic Evaluation of Normalization Approaches in Tandem Mass Tag and Label-Free Protein Quantification Data Using PRONE"

#### Supplementary Methods

##### Spike-In Datasets

*dS1*

The sixth study of the Clinical Proteomic Technology Assessment for Cancer (CPTAC) (1) involves the integration of the Sigma UPS1 standard containing 48 different human proteins spiked at five different concentrations (0.25 fmol UPS1 proteins/µL, 0.74, 2.22, 6.67, and 20 fmol/µL) into a constant protein background of 60 ng/µL Saccharomyces cerevisiae (2). The samples were examined in five different laboratories using seven different instruments. Välikangas *et al.* (3) restricted their study to the replicates examined on the LTQ Orbitrap Velos mass spectrometer at test site 86 and excluded sample group E. The raw files were processed using the Progenesis software and Mascot (4) search engine via Proteome Discoverer (5) for peptide identification. Upon request, the authors provided the resulting protein quantification matrix.

*dS2*

The dS2 dataset consists of nine conditions, each with three replicates. Different concentrations of UPS1 (0.05, 0.125, 0.250, 0.5, 2.5, 5, 12.5, 25, 50 fmol/$\mu$g) were added to a stable yeast lysate background (6). The samples were analyzed using nanoLC-MS/MS on a LTQ-Velos Orbitrap mass spectrometer. The resulting 27 raw files are deposited in the ProteomeXchange (7) repository with the identifier PXD001819. In this study, we used the MaxQuant (8) output file published in the GitHub repository of proteiNorm (9). Due to the high amount of UPS1 proteins with missing values in samples with 0.05, 0.125, 0.250, and 0.5 fmol/$\mu$g of spike-in proteins, we excluded those samples for this study.

*dS3*

Varying amounts of DH5ɑ *E.coli* digest have been added to a large and constant background of human PANC-1 cell digest (10). The experiment included a total of five groups of *E.coli* spike-in, each containing four replicates (percentage of *E.coli* in total proteins): 3 % (onefold, group A), 4.5 % (1.5-fold, group B), 6 % (twofold, group C), 7.5 % (2.5-fold, group D), and 9 % (threefold, group E). The samples were analyzed by a nanoLC-MS system consisting of a Spark Endurance autosampler, an ultra-high-pressure Eksigent Nano-2D Ultra capillary/nanoLC system, and an Orbitrap Fusion mass spectrometer. The raw files of the dataset are deposited in the ProteomeXchange (7) repository with the dataset identifier PXD003881. Sticker *et al.* (11) compared different DE analysis software tools on this benchmark spike-in dataset, and therefore re-evaluated the data using MaxQuant (8). The authors provided the proteinGroups.txt file, consisting of 4912 proteins measured in 20 samples, in their GitHub repository accessible at https://github.com/statOmics/MSqRobSumPaper.

*dS4*

Cox *et al.* (12) designed a proteome benchmark dataset of HeLa and *E.coli* lysates mixed at defined ratios. To an equivalent quantity of the human proteome background (60 µg HeLa cells protein extract), the *E.coli* proteome was prepared in two conditions with a 1:3 ratio (10 µg vs. 30 µg) in triplicates. The samples were analyzed via LC combined with electrospray MS/MS on an LTQ Orbitrap with lock mass calibration. Quantification was done using MaxQuant (8), and the output files are available on the ProteomeXchange (7) repository with the identifier PXD000279.

*dS5*

This is the dataset D3 from PXD013277 PRIDE archive of Zhu *et al.* (13). Samples were prepared by spiking *E.coli* proteins in varying amounts (three replicates at 7.5 µg, four replicates at 15 µg, and three replicates at 45 µg) into a background of MCF-7 human protein extract (70 µg). The samples were labeled with 10-plex TMT, pooled into a single sample, and analyzed in a large-scale fractionation using a Q Exactive HF instrument. Phil Wilmarth reanalyzed the data in his public GitHub repository using the PAW pipeline. The data is available on the GitHub repository, accessible at https://github.com/pwilmart/PXD013277_E-coli_spike-ins_MS2-TMT, under the name edgeR input.txt and is utilized in this study.

*dS6*

O'Connell *et al.* (14) designed a benchmark dataset of spike-in yeast proteins to evaluate the performance of two of the most common data-dependent methods for proteome-wide quantification, isobaric-labeling with TMT and LFQ. Four samples were created by spiking the yeast lysate to 3.3 % of the total protein concentration. Four samples and three samples consist of yeast lysate spiked into human lysate to reach 5 % and 10 % of the total protein concentration, respectively. The samples were analyzed on an Orbitrap Fusion Lumos instrument coupled to a Proxeon EASY-nLC 1200 LC pump. The raw data can be accessed from ProteomeXchange (7) with identifier PXD007683.

Ammar *et al.* (15) processed the LFQ and TMT dataset using MaxQuant (8), and the resulting quantitative protein and peptide intensity tables are available at the following website https://www.bio.ifi.lmu.de/ software/msempire/index.html. In this study, the TMT dataset is utilized since it led to the identification of a larger number of yeast and human proteins compared to the LFQ dataset.

#### Biological Datasets

*dB1*

Li *et al.* (16) conducted a TMT experiment to assess the temporal protein expression changes during osteogenic differentiation of human periodontal ligament stem cells at four different time points (D0, D3, D7, and D14) with triplicate biological replication. The same pooled reference sample was included in each of the three 6-plex TMT sets. LC-MS/MS analysis was performed on a Q Exactive mass spectrometer that was coupled to an Easy-nanoLC. MS/MS spectra were searched using Mascot (4), embedded into Proteome Discoverer (5). Since the quantified data was not published, the raw files were extracted from ProteomeXchange (7) with identifier PXD020908 and re-analyzed with MaxQuant (version 1.6.3.3) (8). Experimental mass spectra were matched to reference proteomes of *Homo sapiens* retrieved from UniProtKB. Trypsin/P was set as protease in specific mode allowing for a maximum of two missed cleavages. Carbamidomethylation of cysteine was set as fixed modification whereas oxidation of methionine and acetylation of protein N-terms were set as variable modifications. Mass tolerances were limited to 26 ppm for full scans in the first search as well as fragment ions and to 4.5 ppm for full scans in the main search. A minimum of two peptides including one unique peptide was necessary for protein inference. FDR for peptide-spectrum-matches, peptides and proteins were controlled to 0.05 by target/decoy evaluation.

*dB2*

Hu *et al.* (17) from Johns Hopkins University employed MS for global proteomic quantification of 83 new high-grade serous ovarian adenocarcinoma samples and 23 relevant non-tumor tissue samples. Tumor and non-tumor samples, along with pooled reference samples and technical replicates of a quality control sample, were labeled using TMT and distributed across 13 10-plex TMT sets. LC-MS/MS analysis was conducted using a Q-Exactive mass spectrometer. The raw proteomics data are available via the Clinical Proteomic Tumor Analysis Consortium (CPTAC) Data Portal (https://pdc.cancer.gov/pdc/study/PDC000110). We employed MaxQuant (version 2.3.1.0) (8) to re-analyze the raw files. Data were processed with default parameters unless otherwise specified, including a peptide and protein false discovery rate of 1% and a minimum peptide length of 7 amino acids. Non-default settings included carbamidomethyl as a fixed modification, LysC and trypsin as enzymes, MS2 reporter ion, and TMT10plex labels for modifications. Peptide identification and quantification were performed against the OpenProt database (release 1.6, September 1, 2020) (18), incorporating RefProts from Uniprot release 2019_03_01. The metadata information of the samples was obtained from the supplementary materials of (17) and matched with the sample IDs of the raw proteomics data. Samples SPL050, SPL039, and SPL047 were removed due to missing metadata. Additionally, quality control samples were excluded due to a high median correlation of the quantified proteins in the nine quality control analyses. Consequently, protein intensities from 119 samples (13 pooled reference samples, 23 non-tumor, and 83 tumor samples) were subjected to normalization in this study.

*dB3*

Vehmas *et al.* (19) investigated the effect of an increased estrogen-to-androgen ratio on liver lipid metabolism in males. AROM+ transgenic mice were utilized as a model as these mice possess an increased estrogen-to-androgen ratio due to the overexpression of the human P450 aromatose enzyme, which converts androgens to estrogens. The authors studied the impact of this hormonal imbalance on the liver proteome and transcriptome, and plasma phospholipid profile. Liver samples of five AROM+ and seven wild-type male mice were analyzed on an LTQ Orbitrap Velos Pro mass spectrometer coupled to an EASY-nLC, and the protein data was searched using Proteome Discoverer (5) and the UniProtKB/Swiss-Prot mouse database. The proteomics MS data and non-normalized quantification file have been deposited to the ProteomeXchange (7) Consortium via the PRIDE partner repository with the dataset identifier PXD002025.

#### Normalization Methods

*Simple sample shifting*

Normalization is achieved within this category by shifting sample data to a specific value. This category encompasses mean and median normalization, two types of global intensity normalization, and median absolute deviation (MAD) normalization. Mean normalization adjusts the scale of the samples to have the same mean value, while median normalization shifts the samples toward the same median value (3,20). Global intensity normalization is used in proteomics to adjust for slight variations in sample loading and labeling efficiency (21). Specifically, it adjusts for variations in the total amount of protein in each sample since an equal amount of protein is typically processed for each sample. In the literature, the method of global intensity normalization can be attained either by calculating a global scaling value as the mean or the median of intensity sums of all proteins across all samples (22,23). Moreover, MAD normalization is a method used to correct for scale or dispersion differences in data, reviewed in (24). The method postulates that the median and the spread of the intensity distribution should be constant across all samples. MAD normalization was performed using the performSMADNormalization function from the R/Biocondcutor-package NormalyzerDE (25). Dressler *et al.* (26) introduced a novel normalization approach grounded on the relationship between technical variation and the actual biological DE of a certain proportion of proteins. The approach involves computing the variance adjusted for expression level and the mean correlation with all other proteins, which aids in ranking the proteins based on their likelihood of being not DE. Normalization is then based only on a selection of n DE proteins, employing the standard median normalization method (NormicsMedian). The normalization parameters are subsequently applied to the full dataset. The algorithm was originally developed in Python but was translated into R for integration into the PRONE package.

*Sample-to-reference*

This category comprises normalization methods based on sample-to-reference transformations, including quantile normalization, linear and local regression normalization methods, and trimmed mean of M-values (TMM). These normalization techniques either use an existing sample as a reference or create a reference sample from available samples to normalize the data. Quantile normalization assumes the data to originate from an identical distribution where most protein intensity signals are unchanged among the samples. The sample distributions are forced to be the same on the quantiles of the samples. First, the values of each protein in each sample are ranked from low to high. Next, the average of the values according to the respective sample ranks is computed, and finally, the protein values are replaced by their average rank value (20). Quantile normalization was performed using the normalize.quantiles function from the R/Biocondcutor-package preprocessCore (27).

Linear regression normalization methods assume that the bias in the data is linearly related to the magnitude of the protein intensity measurement. This normalization technique is performed by fitting a least squares regression model, and the normalized values are calculated using the predicted intensities from the regression equation. We explored three variants of the robust linear regression called Rlr, RlrMA, and RlrMACyc. The Rlr method performs a single reference analysis. The reference sample to which all the other samples are normalized is equivalent to the median intensity of all samples. The RlrMA is similar to Rlr, with the exception that the data are MA transformed before normalization, where A refers to the median sample and M is calculated for each sample as the difference of that sample to A. Finally, in RlrMACyc, MA transformation is performed and the normalization of the samples is done pairwise between two samples, with A being the average of the two samples and M the difference. The process is then performed for all sample pairs, and the cycle is repeated a total of three times, which has been found to be sufficient to achieve convergence between iteration cycles (3).

In contrast to linear regression normalization methods, which assume a linear relationship between the bias in the data and the magnitude of protein intensity, local regression normalization methods assume a nonlinear relationship between these quantities. Local weighted regression is used to fit a curve to the data and estimate intensity-dependent differences between pairs of analyses. Two common variants were already previously applied for the normalization and proteomics data and were implemented in this study. Cyclic loess normalization (LoessCyc) is similar to the RlrMACyc method in that it uses MA-transformed data in a cyclic process, whereas fast loess normalization (LoessF) is similar to the RlrMA method. LoessCyc and LoessF was performed using the normalizeCyclicLoess function of the R/Bioconductor-package limma (28).

TMM, initially developed for RNA-seq data (29), was designed to correct for biases introduced by compositional differences between samples, such as the presence of a few highly abundant genes that can affect the relative expression levels of all other genes, as in (30). TMM normalization adjusts for this kind of bias using the concept of an MA plot and applying a double trimming by M- and A-values. We employed the calcNormFactors function of the R/Bioconductor-package edgeR (31) to calculate the normalization factors.

Notably, the cyclic regression methods, RlrMACyc and LoessCyc, can inherently manage missing values, but if a protein in a specific sample contains even a single missing value, the entire row corresponding to that protein will be composed of MVs post-normalization. This accumulation of missing values is a consequence of the cyclic process and the pairwise normalization approach applied to the samples.

*Variance stabilizing*

VSN, initially developed for microarray experiments (32) and reviewed for proteomics data in (3,9,26), aims to stabilize the variance of the data across different samples and conditions, to adjust the scale of the data, and reduce the impact of outliers. It utilizes a set of parametric transformations and maximum likelihood estimation to make the sample variances independent from their mean intensities and bring the samples onto the same scale. VSN was performed using the normalizeCyclicLoess function of the R/Bioconductor-package limma (28). Furthermore, similar to the NormicsMedian method, Dressler *et al.* (26) introduced NormicsVSN. This method applies the original VSN normalization exclusively to the chosen non-DE proteins, and then utilizes the derived parameters to normalize the entire dataset.

*Model-based*

EigenMS (33) and RobNorm (34) are included in the category: model-based normalization. EigenMS is used for normalizing LC-MS peak intensity measurements to remove technical bias while preserving the original group-level differences in the data. EigenMS is formulated in two steps: (i) it fits an analysis of variance model to estimate the effects of the experimental factors on the data using knowledge about the experimental design, and then (ii) applies singular value decomposition to identify systematic trends contributing to significant variation not explained by the experimental factors of interest. Notably, EigenMS can only handle protein intensity values that have more than one valid value per sample group due to the ANOVA model inside of EigenMS. Proteins that do not fulfill this criterion can not be normalized, and all intensities will be replaced with MVs.

The novel approach RobNorm (34) is a model-based robust normalization method developed for labeled quantitative MS-based proteomics. Unlike other normalization methods, RobNorm does not rely on the assumption that protein intensities are similar across all samples, which makes it more effective when working with protein expression data from heterogeneous samples, such as different tissues. It uses the density-power-weight method to estimate and remove the sample effects and to down-weight outliers. In addition, the developers of RobNorm adapted the one-dimensional robust fitting method, previously described in former work, to the structure of proteomics data in order to obtain a robust estimate of the sample effect.

*Batch effect correction*

While tandem mass tag (TMT) multiplexing offers a robust and accurate approach for quantitative proteomics, it is critical to recognize its potential drawbacks, especially regarding data analysis across various TMT batches (35). Brenes *et al.* (35) indicate that batch effects in TMT significantly impact proteomics data, surpassing the effects of actual biological variation. To correct for such non-biological variations, PRONE currently offers two batch effect correction methods.

IRS, initially delineated by Plubell *et al.* (21) in 2017 and further reviewed in (30), uses the internal reference samples for normalizing protein intensities across different TMT experiments. Specifically, the process involves computing the geometric average intensity for each protein across all reference samples. Subsequently, a scaling factor is calculated for each TMT experiment by dividing the geometric average intensity by the corresponding intensity in its reference sample. Ultimately, the data of each TMT experiment is adjusted by multiplying with this derived scaling factor.
The limma (28) package's removeBatchEffect function, here referred to as limBE, employs a linear model to account for technical batch effects in omics data. A linear model is fitted for each variable given a series of conditions as explanatory variables, including batch and treatment effects. Unlike conventional linear or linear mixed models that estimate both treatment and batch effect simultaneously, limBE specifically removes the batch effect from the original data.

#### Implementation

PRONE is available as an R package on Bioconductor, with the source code accessible at https://github.com/daisybio/PRONE. A graphical interface is deployed at https://exbio.wzw.tum.de/prone, with the source code available at (https://github.com/daisybio/PRONE.Shiny.

To simplify the analysis and ensure a compact data structure across the workflow, the functions are based upon a SummarizedExperiment container (36). This ensures easy handling of the data during the six-step workflow of PRONE (**Figure 2**). In addition to the protein intensity table and the metadata information, other additional columns originating from MaxQuant (8), for instance, are allowed and can be accessed from the SummarizedExperiment object at any time. Furthermore, data derived from the different normalization techniques is stored as assays within the SummarizedExperiment, streamlining the programming process by requiring only a single object instead of multiple objects for each individual normalization technique. Finally, DE analysis can be performed simultaneously across all normalization methods stored within the SummarizedExperiment container. This results in a single data table that consolidates the DE outcomes for all specified pairwise comparisons and normalization methods. Consequently, the implementation of PRONE streamlines data management and storage, enhancing ease of use for the user. This configuration of PRONE allows users to readily develop new visualization functions and integrate other normalization techniques, thereby enhancing the comprehensiveness of the package.

### Supplementary Tables

***Supplementary Table 1: Overview table of preprocessing of spike-in datasets.*** *Initially, the datasets were preprocessed by discarding proteins that exhibited missing values (MVs) in all samples, and, if applicable, were only identifiable with a modification site, were reverse hits, or potential contaminants. Since the accuracy of MS-based proteomics data is challenged by the presence of MVs, the rate of MVs was analyzed, and a filtering threshold of 80% was applied to all spike-in datasets. This threshold value denotes the minimal percentage of samples necessitating a valid value for a given protein. If this criterion was not fulfilled, the specific protein was eliminated from the dataset. The final number of proteins per dataset is provided in the last column with the number of spike-in proteins being shown in brackets. POMA outlier detection and manual evaluation of sample outliers were performed, and no outlier samples were detected.*

| Dataset | Number of Samples | Number of Conditions | Initial Number of Proteins | Number of Proteins after Pre-Filtering | MV Rate | Final Number of Proteins |
|---|---|---|---|---|---|---|
| dS1 | 12 | 4 | 736 | 736 | 0.6 % | 731 (36) |
| dS2 | 15 | 5 | 1092 | 1009 | 23.48 % | 656 (20) |
| dS3 | 20 | 5 | 4912 | 4712 | 7.77 % | 4112 (602) |
| dS4 | 6 | 2 | 6694 | 6546 | 18.33 % | 4951 (1530) |
| dS5 | 10 | 3 | 9650 | 9650 | 0 % | 9650 (2091) |
| dS6 | 11 | 3 | 8414 | 8346 | 0.04 % | 8341 (1210) |

***Supplementary Table 2: Pooled median absolute deviation of spike-in datasets.*** *The pooled median absolute deviation (PMAD) was calculated across all sample groups within a specific dataset. Here, the PMAD values are summarized across sample groups by showing the median and the median absolute deviation in brackets. Normalization methods and log2-transformation are sorted by median PMAD over all spike-in datasets. A higher PMAD indicates higher intragroup variation. High and low values for PMAD are highlighted in red and blue, respectively.*

| Normalization | dS1 | dS2 | dS3 | dS4 | dS5 | dS6 |
|---|---|---|---|---|---|---|
| MAD | 0.088 (0.005) | 0.084 (0.008) | 0.082 (0.011) | 0.064 (0.006) | 0.024 (0.004) | 0.016 (0.001) |
| EigenMS | 0.273 (0.048) | 0.09 (0.08) | 0.155 (0.005) | 0.125 (0.03) | 0.048 (0.001) | 0.031 (0.002) |
| LoessCyc | 0.273 (0.015) | 0.195 (0.026) | 0.179 (0.019) | 0.183 (0.011) | 0.073 (0.016) | 0.045 (0.002) |
| RlrMACyc | 0.254 (0.03) | 0.189 (0.006) | 0.177 (0.026) | 0.183 (0.01) | 0.071 (0.006) | 0.044 (0.001) |
| VSN | 0.195 (0.013) | 0.207 (0.018) | 0.199 (0.024) | 0.199 (0.018) | 0.067 (0.009) | 0.045 (0) |
| RLR | 0.269 (0.033) | 0.206 (0.043) | 0.199 (0.022) | 0.196 (0.012) | 0.068 (0.01) | 0.045 (0.001) |
| RlrMA | 0.268 (0.034) | 0.207 (0.03) | 0.199 (0.024) | 0.196 (0.016) | 0.068 (0.01) | 0.045 (0.001) |
| LoessF | 0.265 (0.035) | 0.208 (0.033) | 0.199 (0.024) | 0.195 (0.015) | 0.069 (0.008) | 0.045 (0) |
| RobNorm | 0.27 (0.034) | 0.207 (0.018) | 0.199 (0.024) | 0.198 (0.016) | 0.068 (0.01) | 0.046 (0.001) |
| Median | 0.277 (0.033) | 0.216 (0.013) | 0.2 (0.027) | 0.199 (0.018) | 0.068 (0.011) | 0.046 (0.002) |
| Quantile | 0.289 (0.025) | 0.21 (0.041) | 0.2 (0.025) | 0.199 (0.021) | 0.068 (0.01) | 0.045 (0.001) |
| NormicsVSN | 0.273 (0.036) | 0.208 (0.016) | 0.202 (0.023) | 0.202 (0.021) | 0.068 (0.011) | 0.046 (0.002) |
| GlobalMean | 0.277 (0.042) | 0.21 (0.028) | 0.201 (0.026) | 0.203 (0.019) | 0.068 (0.01) | 0.049 (0.004) |
| GlobalMedian | 0.277 (0.042) | 0.21 (0.028) | 0.201 (0.026) | 0.203 (0.019) | 0.068 (0.01) | 0.049 (0.004) |
| Mean | 0.277 (0.042) | 0.211 (0.027) | 0.201 (0.026) | 0.208 (0.026) | 0.068 (0.01) | 0.049 (0.004) |
| NormicsMedian | 0.276 (0.043) | 0.22 (0.017) | 0.206 (0.025) | 0.201 (0.02) | 0.072 (0.004) | 0.047 (0.002) |
| log2 | 0.341 (0.061) | 0.22 (0.028) | 0.241 (0.051) | 0.199 (0.014) | 0.083 (0.004) | 0.052 (0) |
| TMM | 0.323 (0.042) | 0.229 (0.031) | 0.245 (0.064) | 0.199 (0.018) | 0.087 (0.002) | 0.053 (0.008) |

***Supplementary Table 3: Pearson correlation of spike-In datasets.*** *The Pearson correlation was calculated across all sample groups within a specific dataset. Here, the correlation values are summarized across sample groups by showing the median and the median absolute deviation in brackets. Normalization methods and log2-transformation are sorted by median Pearson correlation over all spike-in datasets. A high Pearson correlation indicates high intragroup similarity. High and low values of Pearson correlation are highlighted in blue and red, respectively. For dS2, dS3, dS5 and dS6, only the highest value of the intragroup Pearson correlation is highlighted.*

| **Normalization** | **dS1** | **dS2** | **dS3** | **dS4** | **dS5** | **dS6** |
|---|---|---|---|---|---|---|
| EigenMS | 0.968 (0.031) | 0.999 (0.001) | 0.986 (0.003) | 0.995 (0.004) | 0.999 (0) | 1 (0) |
| RlrMACyc | 0.959 (0.038) | 0.963 (0.009) | 0.977 (0.004) | 0.988 (0.001) | 0.997 (0.001) | 0.999 (0) |
| LoessCyc | 0.955 (0.034) | 0.962 (0.009) | 0.977 (0.003) | 0.988 (0.001) | 0.997 (0.001) | 0.999 (0) |
| VSN | 0.981 (0.018) | 0.962 (0.009) | 0.974 (0.003) | 0.987 (0.001) | 0.997 (0.001) | 0.999 (0) |
| LoessF | 0.956 (0.03) | 0.962 (0.008) | 0.975 (0.003) | 0.987 (0.001) | 0.997 (0.001) | 0.999 (0) |
| Quantile | 0.955 (0.031) | 0.963 (0.009) | 0.974 (0.003) | 0.988 (0.001) | 0.997 (0.001) | 0.999 (0) |
| Mean | 0.951 (0.032) | 0.962 (0.009) | 0.974 (0.003) | 0.987 (0.001) | 0.997 (0.001) | 0.999 (0) |
| NormicsVSN | 0.952 (0.032) | 0.962 (0.009) | 0.974 (0.003) | 0.987 (0.001) | 0.997 (0.001) | 0.999 (0) |
| log2 | 0.951 (0.032) | 0.962 (0.009) | 0.974 (0.003) | 0.987 (0.001) | 0.997 (0.001) | 0.999 (0) |
| NormicsMedian | 0.951 (0.032) | 0.962 (0.009) | 0.974 (0.003) | 0.987 (0.001) | 0.997 (0.001) | 0.999 (0) |
| Median | 0.951 (0.032) | 0.962 (0.009) | 0.974 (0.003) | 0.987 (0.001) | 0.997 (0.001) | 0.999 (0) |
| RobNorm | 0.951 (0.032) | 0.962 (0.009) | 0.974 (0.003) | 0.987 (0.001) | 0.997 (0.001) | 0.999 (0) |
| TMM | 0.951 (0.032) | 0.962 (0.009) | 0.974 (0.003) | 0.987 (0.001) | 0.997 (0.001) | 0.999 (0) |
| MAD | 0.951 (0.032) | 0.962 (0.009) | 0.974 (0.003) | 0.987 (0.001) | 0.997 (0.001) | 0.999 (0) |
| GlobalMedian | 0.951 (0.032) | 0.962 (0.009) | 0.974 (0.003) | 0.987 (0.001) | 0.997 (0.001) | 0.999 (0) |
| GlobalMean | 0.951 (0.032) | 0.962 (0.009) | 0.974 (0.003) | 0.987 (0.001) | 0.997 (0.001) | 0.999 (0) |
| RLR | 0.951 (0.032) | 0.962 (0.009) | 0.974 (0.003) | 0.987 (0.001) | 0.997 (0.001) | 0.999 (0) |
| RlrMA | 0.95 (0.031) | 0.962 (0.008) | 0.974 (0.003) | 0.988 (0.001) | 0.997 (0.001) | 0.999 (0) |

***Supplementary Table 4: Overview table of preprocessing of biological datasets.*** *Two TMT datasets and one LFQ dataset were employed in this study. Initially, the datasets were preprocessed by discarding proteins that exhibited missing values (MVs) in all samples, were only identifiable with a modification site, were reverse hits, or potential contaminants. Since the accuracy of MS-based proteomics data is challenged by the presence of MVs, the rate of MVs was analyzed, and a filtering threshold of 80% was applied to all biological datasets. This threshold value denotes the minimal percentage of samples necessitating a valid value for a given protein. If this criterion was not fulfilled, the specific protein was eliminated from the dataset. Finally, the outlier detection method of POMA was applied to detect outlier samples; however, no samples were excluded from the analysis.*

| Dataset | LFQ or TMT | Number of Samples | Number of Conditions | Initial Number of Proteins | Number of Proteins after Pre-Filtering | MV Rate | Final Number of Proteins | Samples removed with POMA |
|---|---|---|---|---|---|---|---|---|
| dB1 | TMT | 12 (+ 4 refs) | 4 | 6200 | 6200 | 10,71% | 4848 | 0 |
| dB2 | TMT | 106 (+ 13 refs) | 2 | 10564 | 10203 | 18,28% | 7207 | 4 |
| dB3 | LFQ | 12 | 2 | 1499 | 1451 | 0,09% | 1450 | 0 |

#### Supplementary Figures

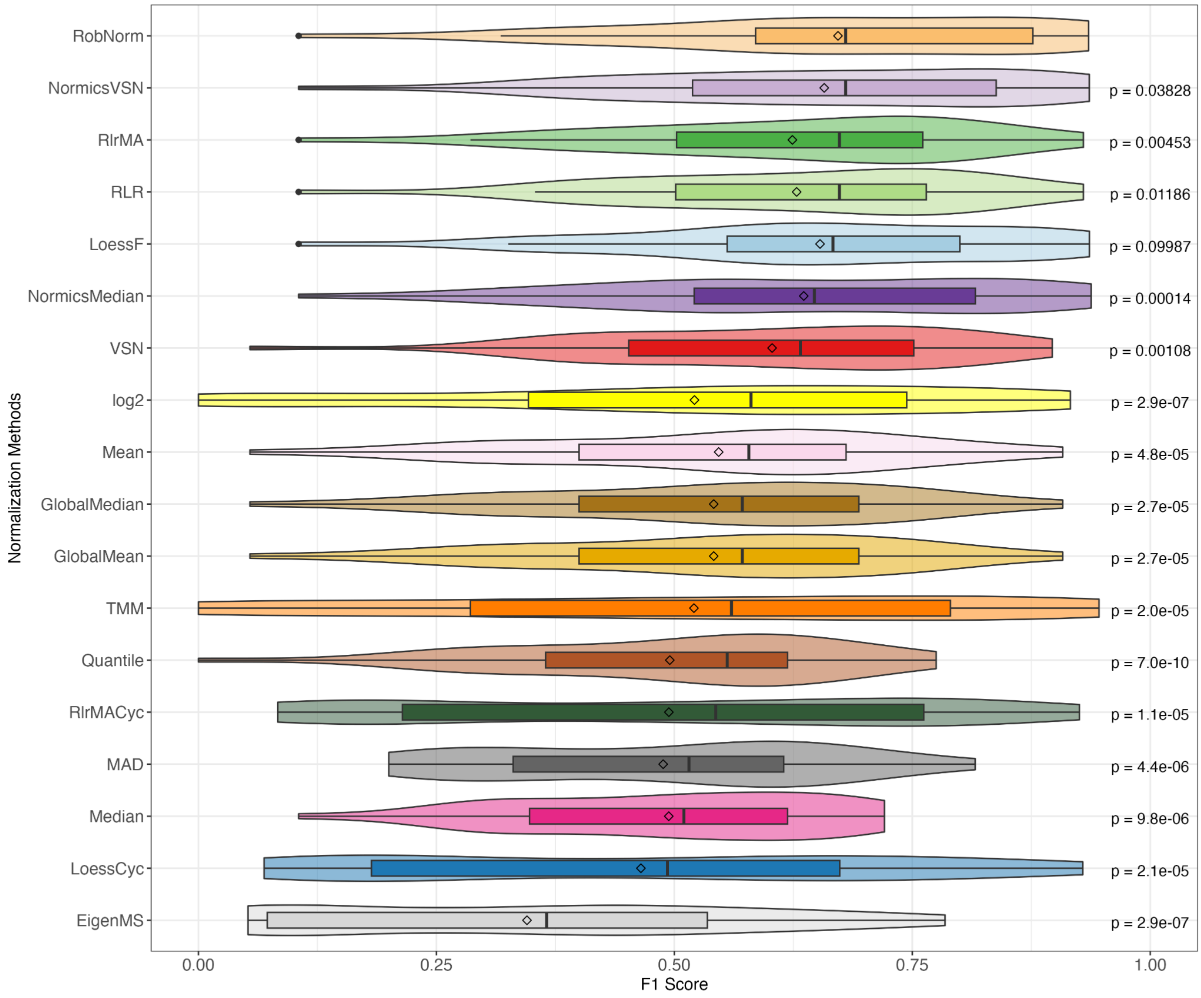


***Supplementary Figure 1: Distributions of F1 scores across spike-in datasets.*** *The F1 scores of all normalization methods across all pairwise comparisons in all spike-in datasets were summarized by using violin plots combined with boxplots. Paired Wilcoxon rank sum tests were conducted to compare the F1 scores of RobNorm with those of all other normalization techniques.*

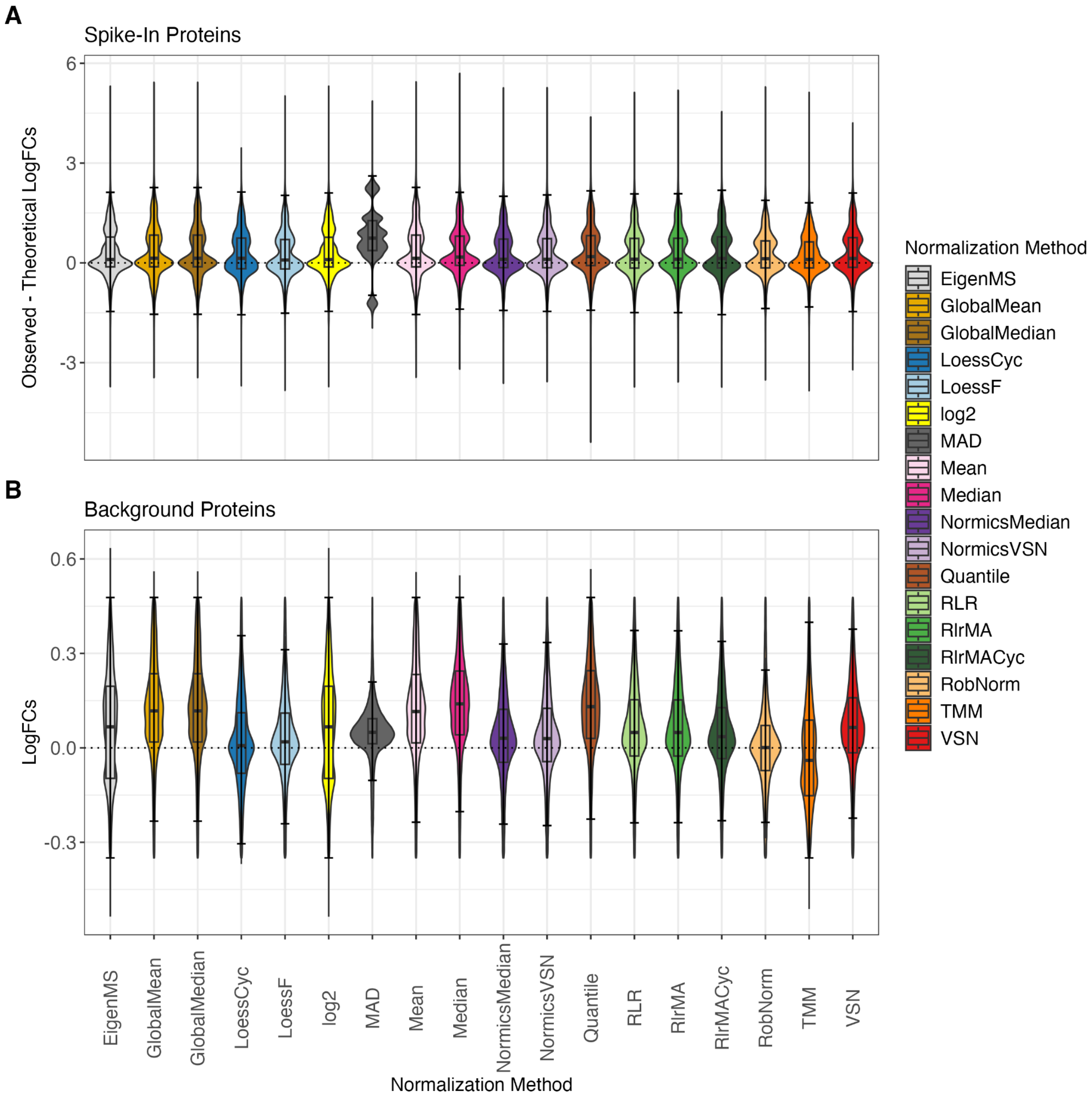


***Supplementary Figure 2: Effect of normalization on log fold changes.*** DE analyses were performed with limma for all pairwise comparisons of all spike-in datasets. *The log fold changes (logFCs) of spike-in proteins were compared to theoretical logFCs by calculating the difference between the observed and theoretical values (A). The logFCs of the background proteins are expected to be centered around 0 since their concentrations remain constant across the sample groups (B). In (B), violin plots visualize all logFC values without outliers, i.e., those values that are within the 1.5-interquartile range (whiskers). In contrast, boxplots are based on all logFC values, including the calculation of the median.*

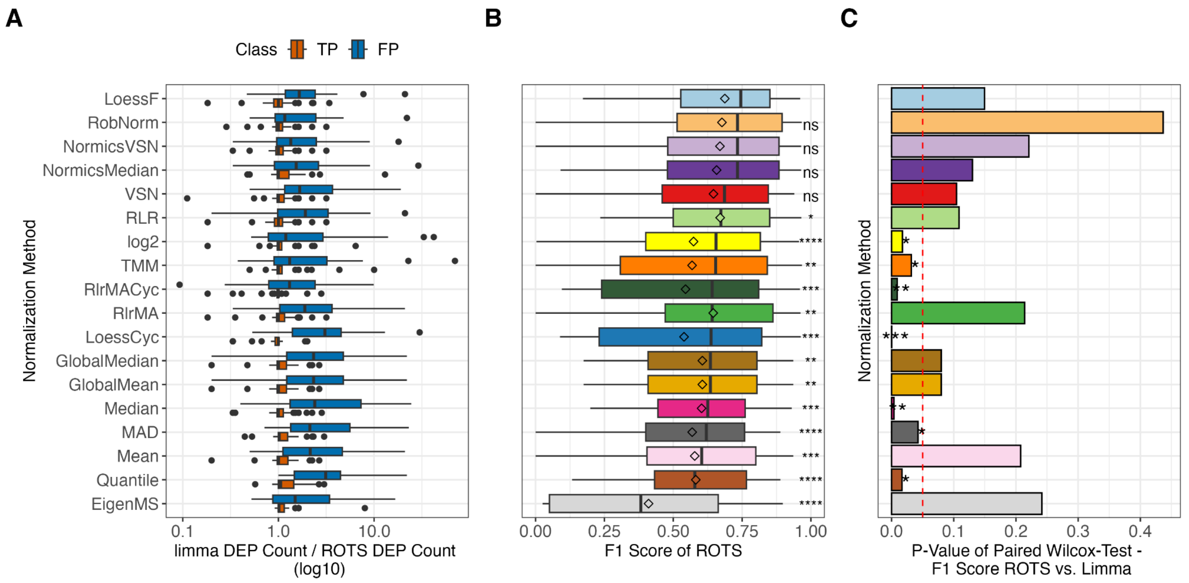


***Supplementary Figure 3: Influence of the differential expression analysis method on the detection of true and false positives in spike-in datasets.*** *The number of true positives (TPs) and false positives (FPs), generated by differential expression analyses using limma divided by those produced by ROTS (A). Ratios exceeding 1 indicate a higher number of proteins identified by limma compared to ROTS. F1 scores (B) were computed based on the DE results of ROTS for all normalization methods, with the normalization methods being sorted according to the median F1 scores across all comparisons and spike-in datasets. Paired Wilcoxon rank sum tests were conducted to compare the F1 scores of LoessF with those of all other normalization techniques. Furthermore, F1 scores of all pairwise comparisons across all spike-in datasets of limma and ROTS were compared in a paired Wilcoxon rank sum test per normalization method with p-values per normalization method being reported in (C). The red dashed line indicates the threshold for statistical significance (p-value < 0.05). *p ≤ 0.05, **p ≤ 0.01, ***p ≤ 0.001, **** p ≤ 0.0001.*

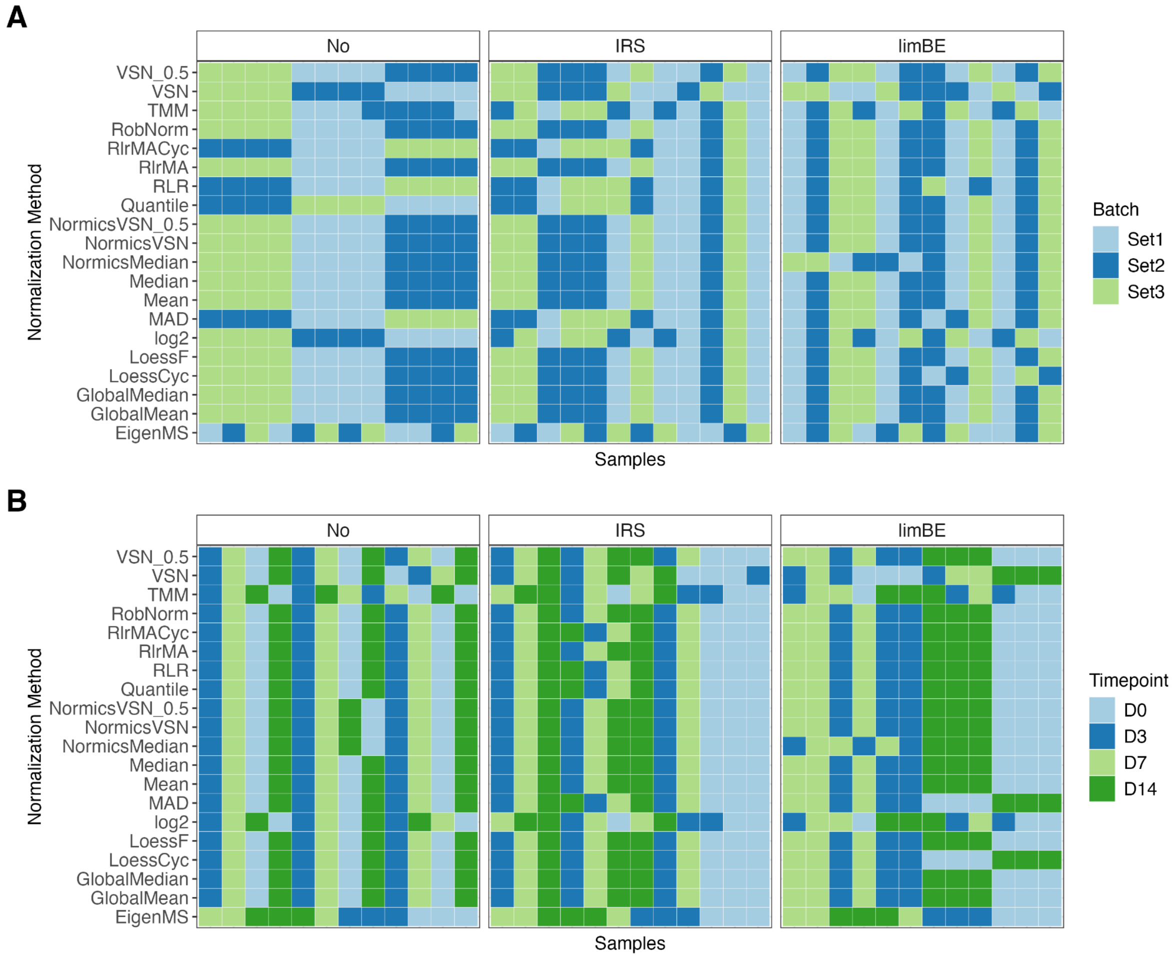


***Supplementary Figure 4: Batch effect assessment of the cell culture dataset dB1 using hierarchical clustering per normalization method.*** *Samples of data normalized using a specific normalization method without batch effect correction (No), with internal reference scaling (IRS), or limma::removeBatchEffects (limBE), were hierarchically clustered and the samples were colored according to TMT-batch (A) or condition (B), respectively. We anticipate that, following batch effect correction, the samples will primarily cluster based on biological condition rather than TMT batch.*

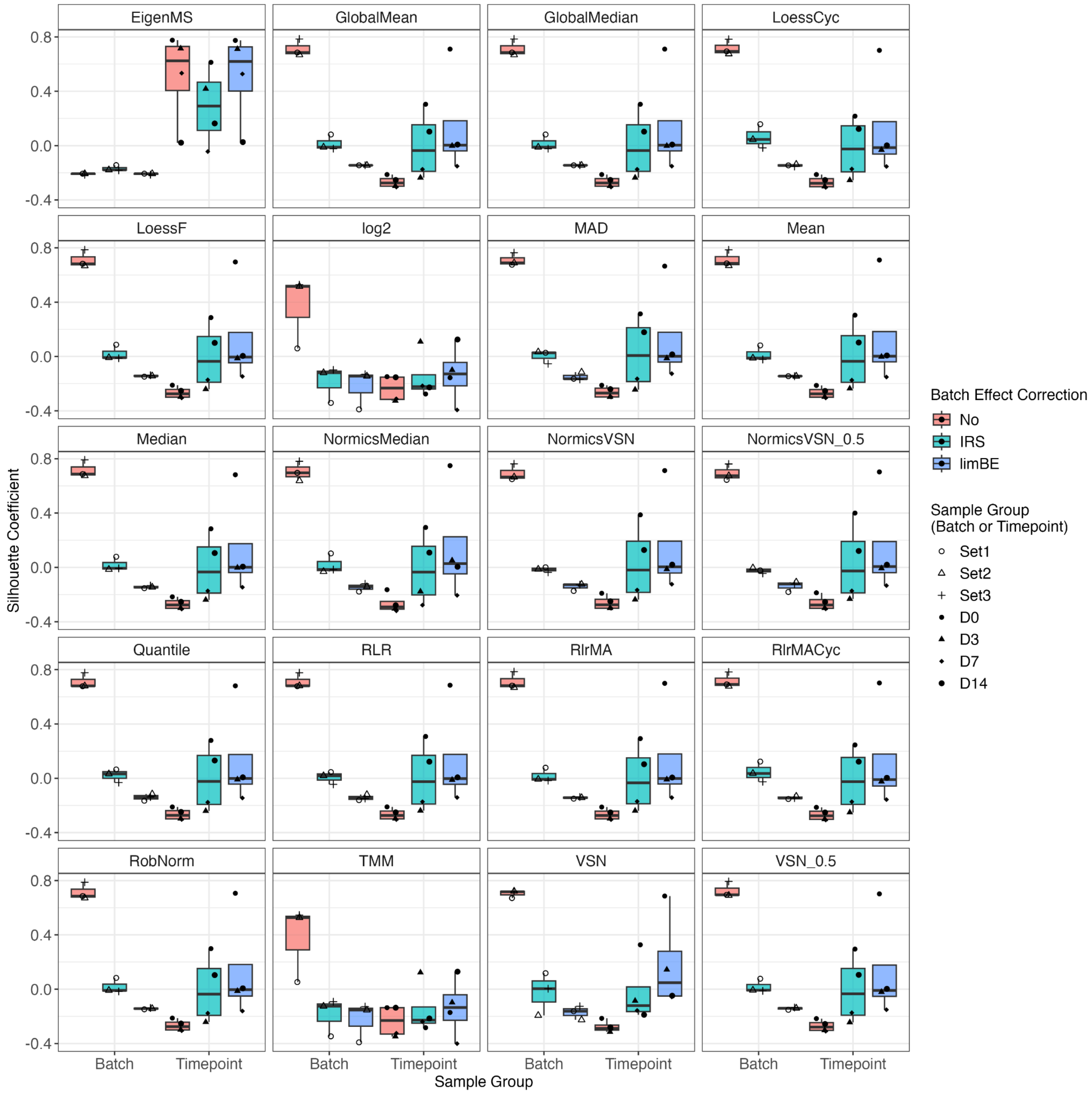


***Supplementary Figure 5: Batch effect assessment of the cell culture dataset dB1 using Silhouette coefficients per normalization method.*** *Euclidean distances between samples of the cell culture dataset dB1 were computed based on the first three principal components. The Silhouette coefficient was calculated for each sample group, batch- and time point-wise, across all combinations of normalization technique and batch effect correction techniques, to quantify how well each sample fits within its assigned group.*

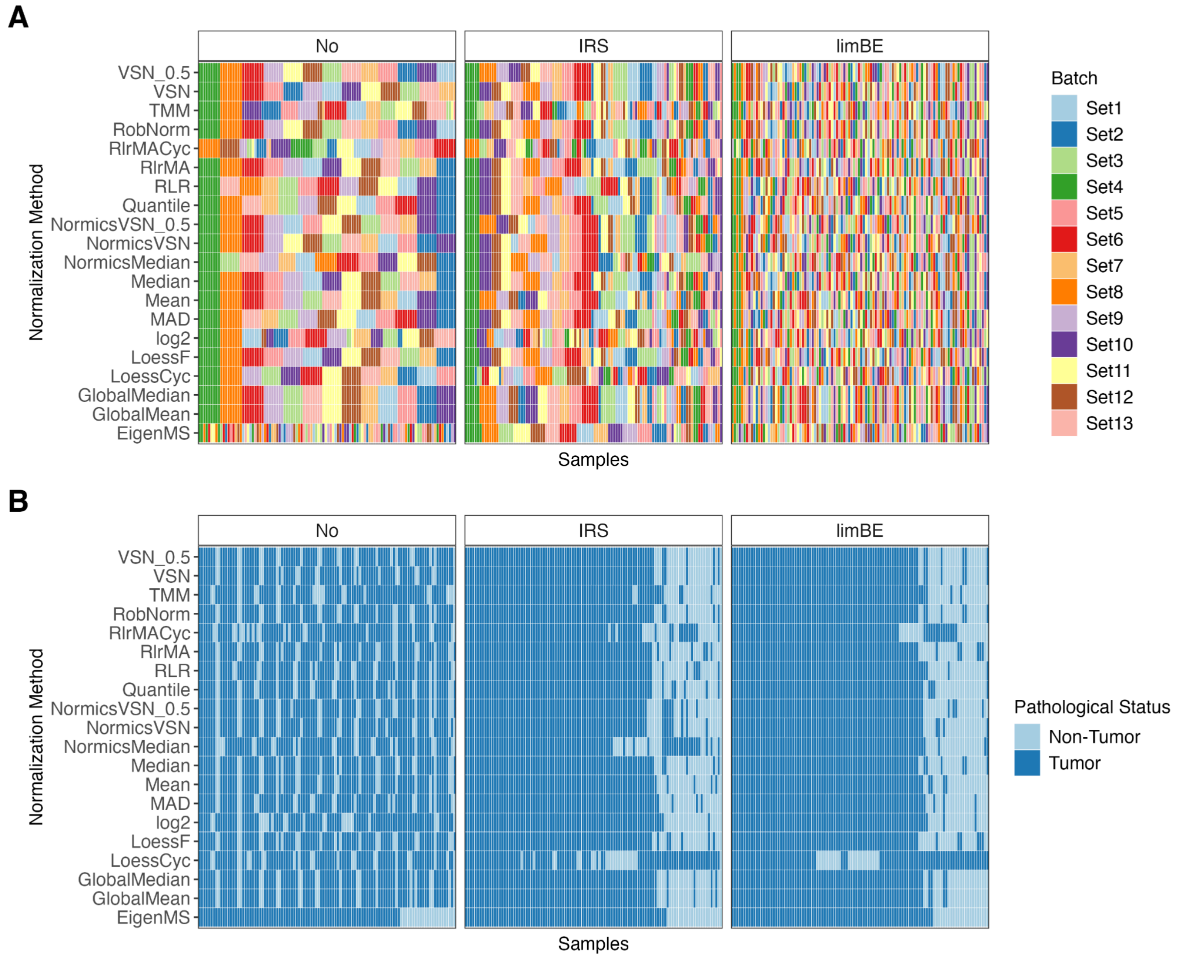


***Supplementary Figure 6: Batch effect assessment of the clinical cancer dataset dB2 using hierarchical clustering per normalization method.*** *Samples of data normalized using a specific normalization method without batch effect correction (No), with internal reference scaling (IRS), or limma::removeBatchEffects (limBE), were hierarchically clustered and the samples were colored according to TMT-batch (A) or condition (B), respectively. We anticipate that, following batch effect correction, the samples will primarily cluster based on biological condition rather than TMT batch.*

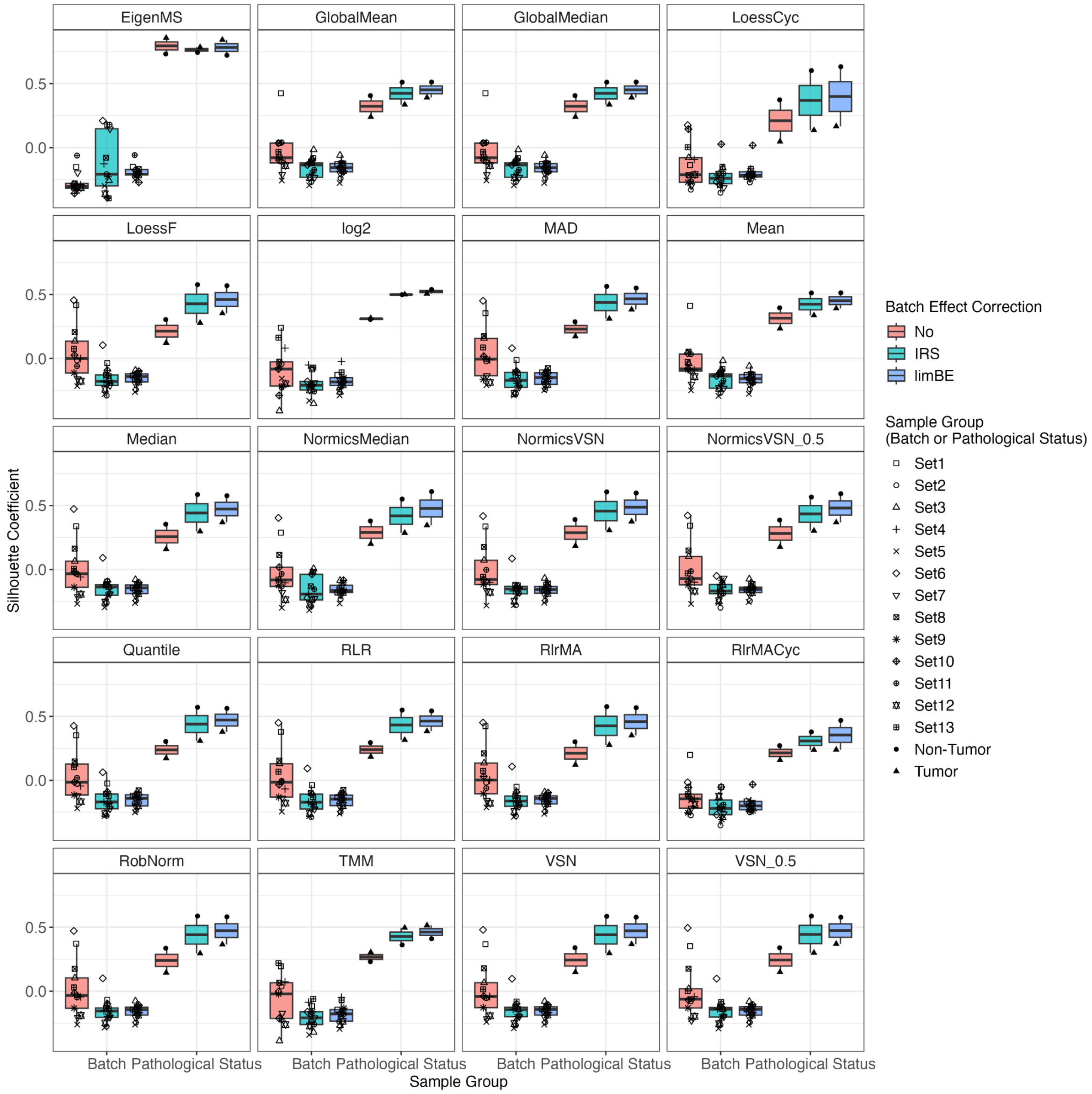


***Supplementary Figure 7: Batch effect assessment of the clinical cancer dataset dB2 using Silhouette coefficients per normalization method.*** *Euclidean distances between samples of the clinical cancer dataset dB2 were computed based on the first three principal components. The Silhouette coefficient was calculated for each sample group, batch- and pathological status-wise, across all combinations of normalization technique and batch effect correction techniques, to quantify how well each sample fits within its assigned group.*

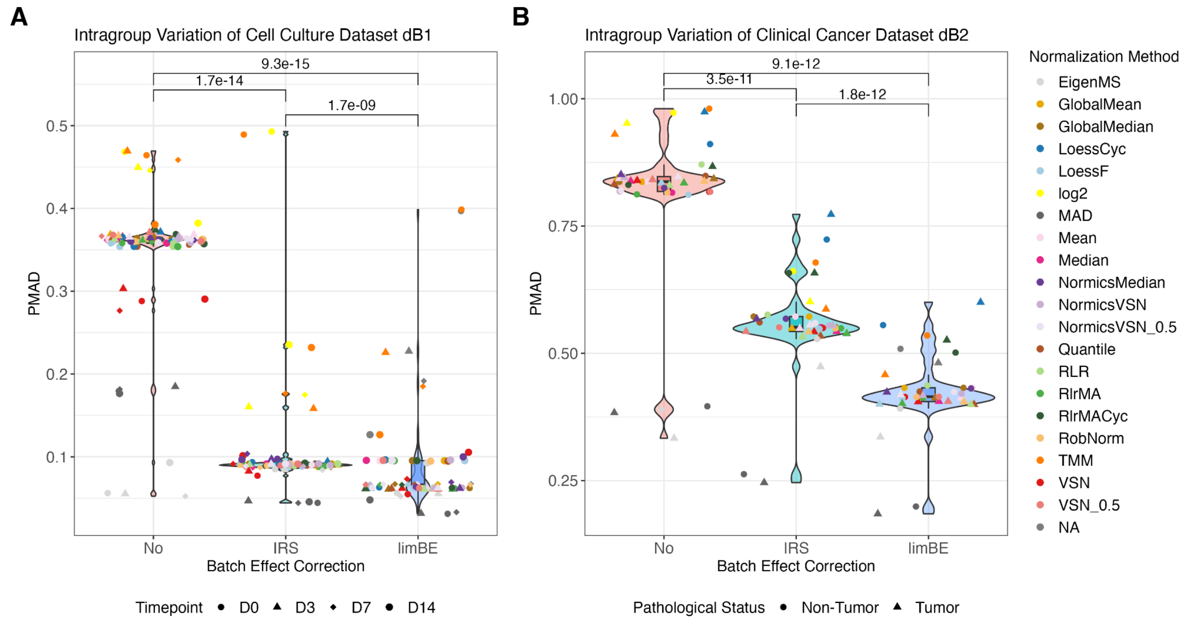


***Supplementary Figure 8: Violin/box plots of intragroup variation for two biological TMT datasets.*** *Pooled median absolute deviation (PMAD) was calculated for each normalization method across samples per condition on data without batch effect correction (No), with internal reference scaling (IRS), and with limma::removeBatchEffects (limBE) on the cell culture dataset dB1 (A) with samples originating from four different time points (D0, D3, D7, D14) and the clinical cancer dataset dB2 (B) with non-tumor and tumor samples. A paired Wilcoxon rank sum test was performed to assess the impact of the batch effect correction on intragroup variation.*

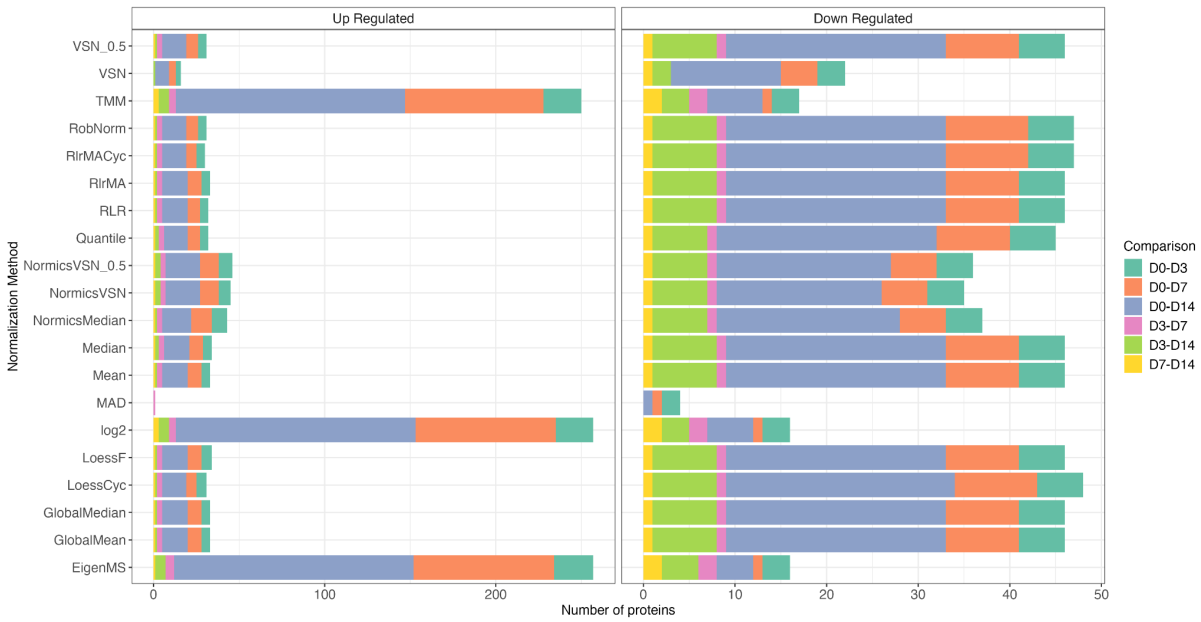


***Supplementary Figure 9: Differential expression results of all pairwise comparisons of the cell culture dataset dB1.*** *Bar plots showing the number of up- and down-regulated differentially expressed proteins of all pairwise comparisons for each normalization method with limBE applied on*

*top to remove batch effects of the cell culture dataset dB1. DE proteins are classified as up- and down-regulated using |logFC| > 1 and Benjamini-Hochberg adjusted p-value < 0.05 was applied.*

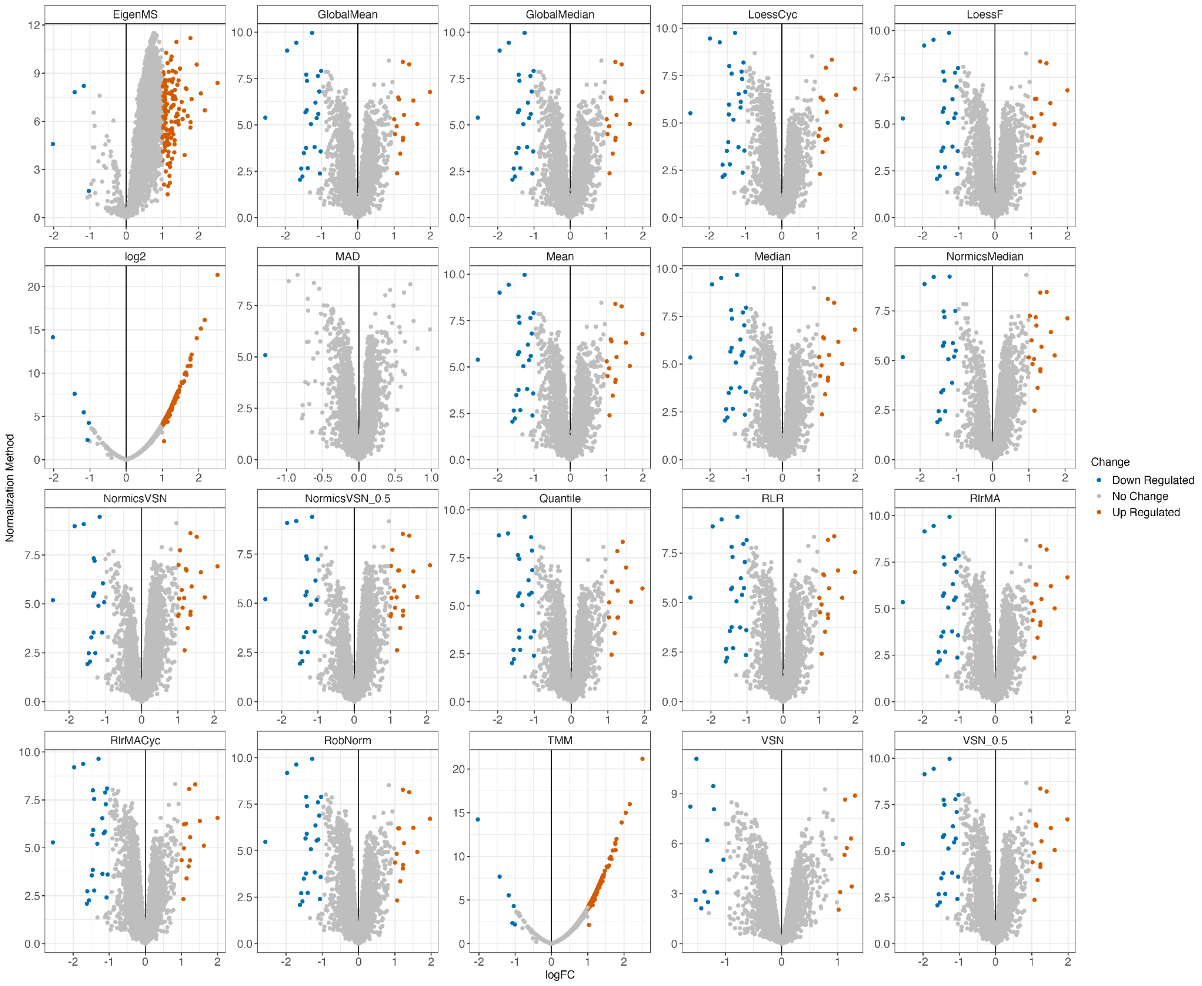


***Supplementary Figure 10: Volcano plots of comparison D0-D14 of cell culture dataset dB1.*** *Individual volcano plots per normalization technique with limBE applied on top to remove batch effects of DE results of the pairwise comparison D0-D14 of the cell culture dataset dB1. DE proteins are classified as up- and down-regulated using |logFC| > 1 and Benjamini-Hochberg adjusted p-value < 0.05 was applied.*

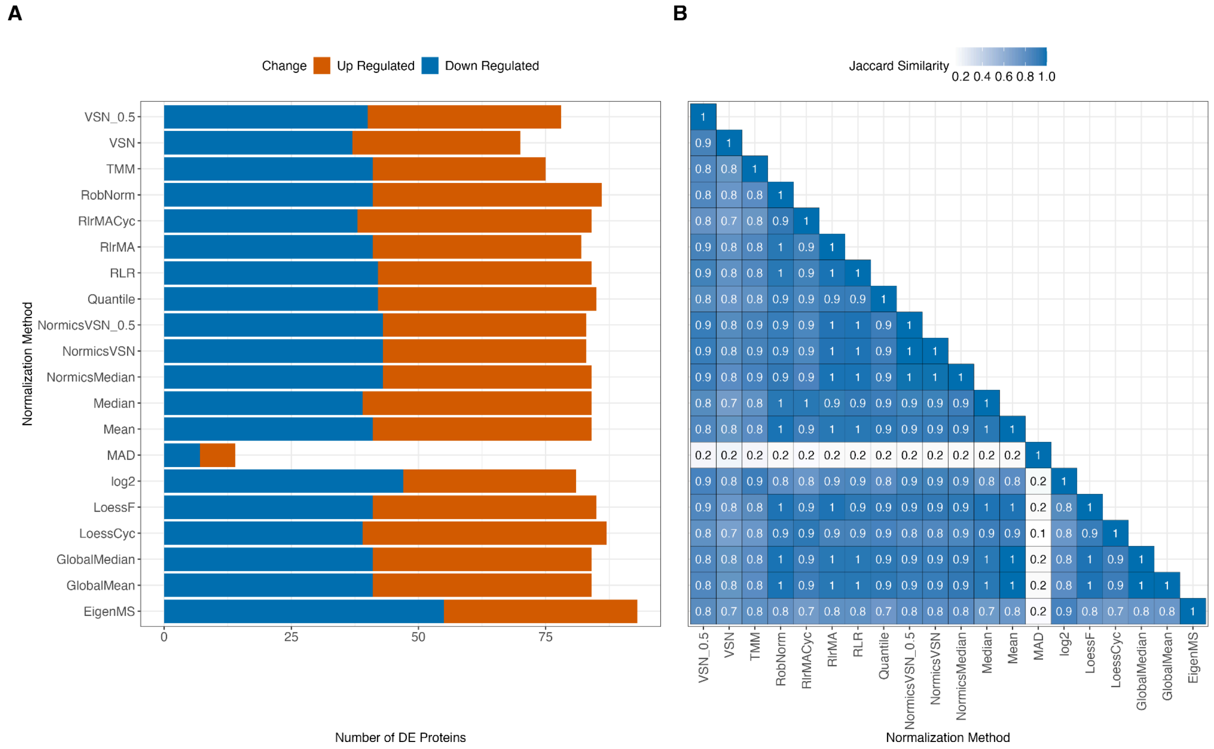


***Supplementary Figure 11: Differential expression results of the LFQ dataset dB3.*** *(A) A barplot showing the number of differentially expressed (DE) proteins of the pairwise comparison for each normalization method in the LFQ dataset dB3 (A). DE proteins were classified as up- and down-regulated using |logFC| > 1 and Benjamini-Hochberg adjusted p-value < 0.05 was applied. (B) A heatmap of the Jaccard similarity coefficients between all normalization techniques based on the DE results of (A).*

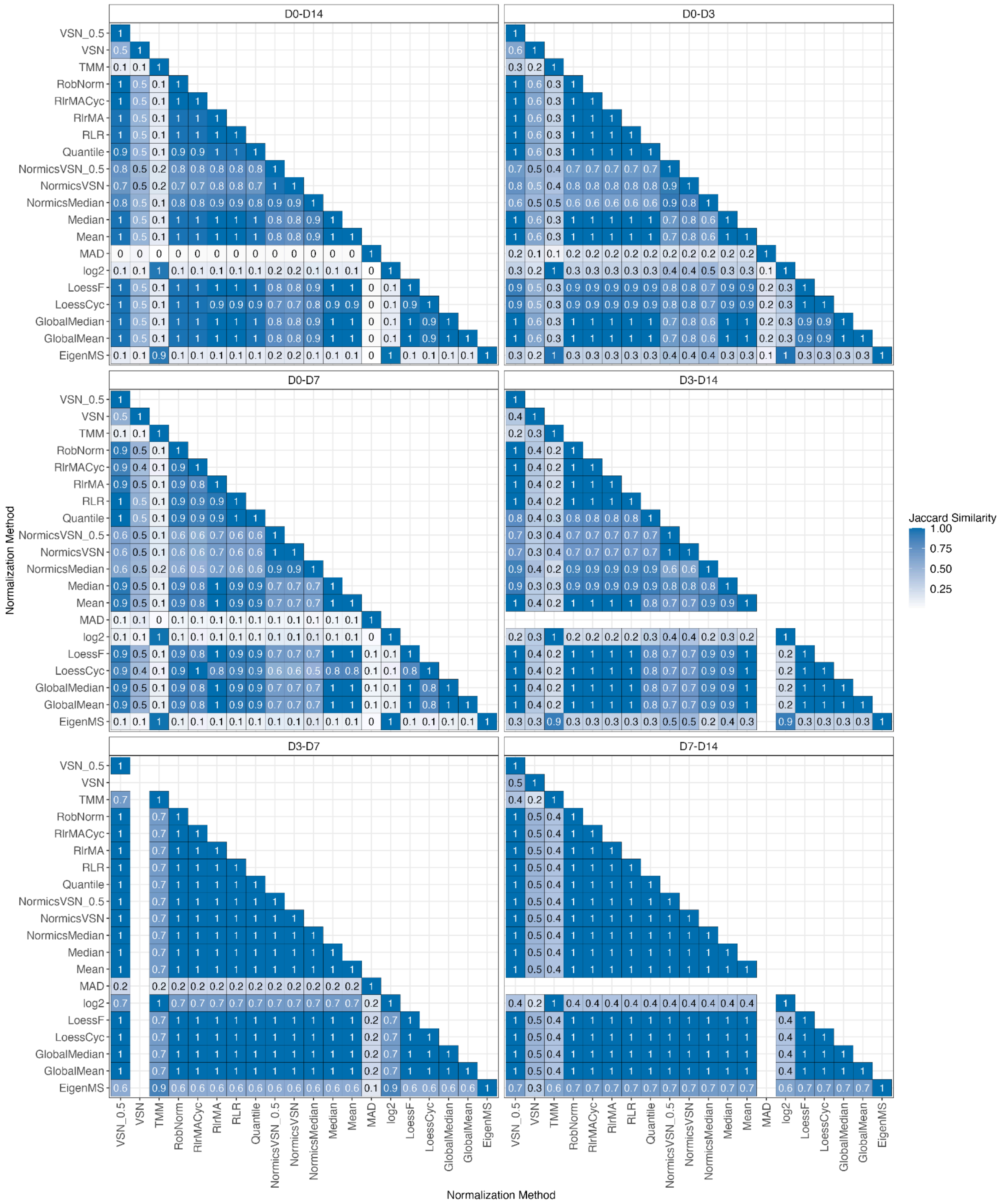


***Supplementary Figure 12: Jaccard similarity coefficients of all pairwise timepoint comparisons of the cell culture dataset dB1.*** *Heatmaps displaying Jaccard similarity coefficients between all pairs of normalization methods with limBE applied on top to remove batch effects based on the DE results of (Supplementary Figure 8) for cell culture dataset dB1.*

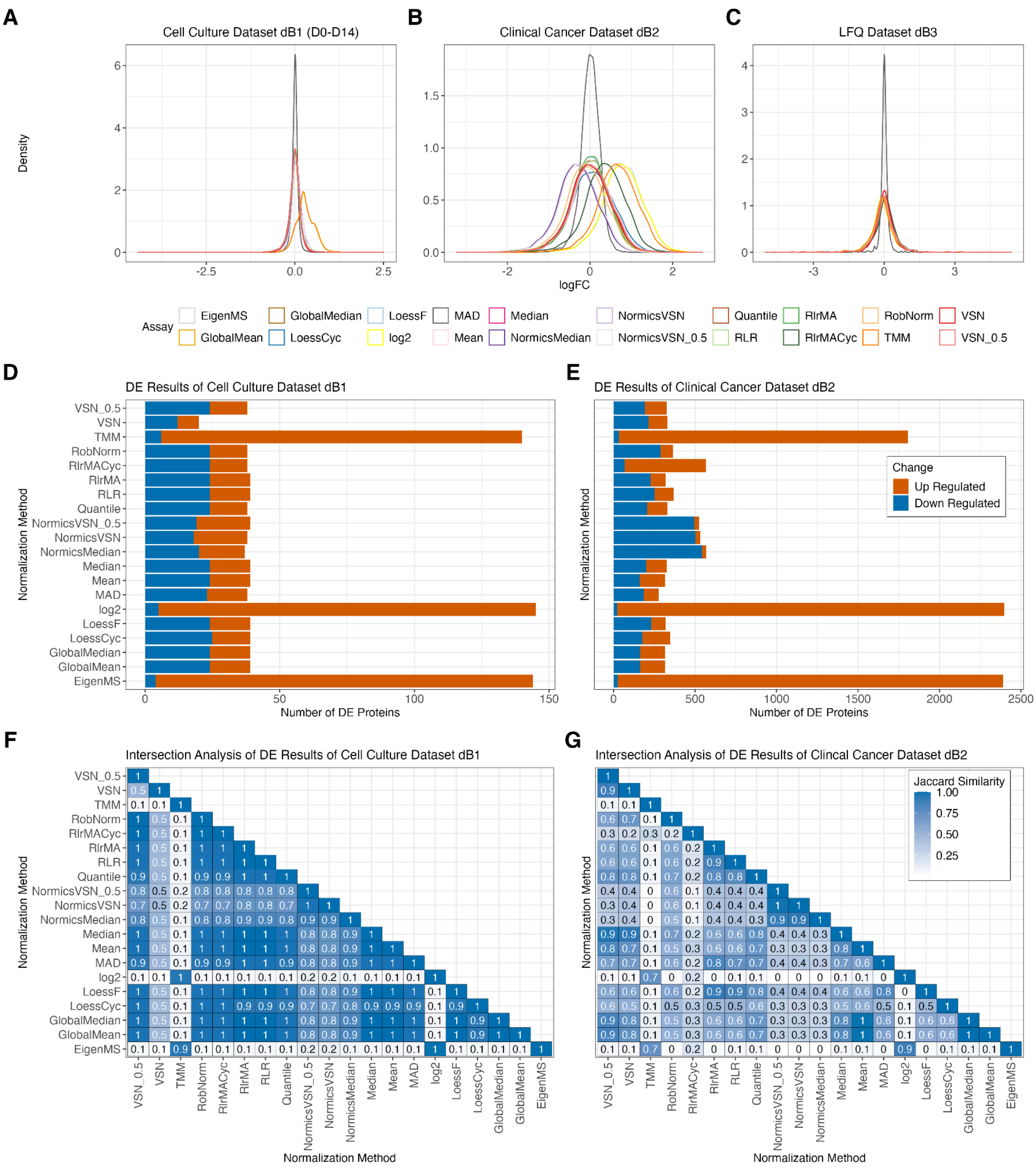


***Supplementary Figure 13: Differential expression results of biological datasets with adjusted logFC threshold.*** *Log fold changes (logFC) of differential expression (DE) results from different normalization methods are shown for the cell culture dataset dB1 (A), the clinical dataset dB2 (B), and the LFQ dataset dB3 (C). Subpanels D-G follow the same structure as Figure 6, but with a less stringent logFC threshold of 0.5 applied for MAD.*
